# Resolving genetic linkage reveals patterns of selection in HIV-1 evolution

**DOI:** 10.1101/711861

**Authors:** Muhammad S. Sohail, Raymond H. Y. Louie, Matthew R. McKay, John P. Barton

## Abstract

Identifying the genetic drivers of adaptation is a necessary step in understanding the dynamics of rapidly evolving pathogens and cancer. However, signals of selection are obscured by the complex, stochastic nature of evolution. Pervasive effects of genetic linkage, including genetic hitchhiking and clonal interference between beneficial mutants, challenge our ability to distinguish the selective effect of individual mutations. Here we describe a method to infer selection from genetic time series data that systematically resolves the confounding effects of genetic linkage. We applied our method to investigate patterns of selection in intrahost human immunodeficiency virus (HIV)-1 evolution, including a case in an individual who develops broadly neutralizing antibodies (bnAbs). Most variants that arise are observed to have negligible effects on inferred selection at other sites, but a small minority of highly influential variants have strong and far-reaching effects. In particular, we found that accounting for linkage is crucial for estimating selection due to clonal interference between escape mutants and other variants that sweep rapidly through the population. We observed only modest selection for antibody escape, in contrast with strong selection for escape from CD8+ T cell responses. Weak selection for escape from antibody responses may facilitate bnAb development by diversifying the viral population. Our results provide a quantitative description of the evolution of HIV-1 in response to host immunity, including selection on the viral population that accompanies bnAb development. More broadly, our analysis argues for the importance of resolving linkage effects in studies of natural selection.

---

Evolving populations exhibit complex dynamics. Cancers (1–6) and pathogens such as HIV-1 (7–9) and influenza (10, 11) generate multiple beneficial mutations that increase fitness or allow them to escape immunity. Subpopulations with different beneficial mutations then compete with one another for dominance, referred to as clonal interference, resulting in the loss of some mutations that increase fitness (12). Neutral or deleterious mutations can also hitchhike to high frequencies if they occur on advantageous genetic backgrounds (13). Experiments have demonstrated that these features of genetic linkage are pervasive in nature (14–16).

Linkage makes distinguishing the fitness effects of individual mutations challenging because their dynamics are contingent on the genetic background on which they appear. Lineage tracking experiments can be used to identify beneficial mutations (17), but they cannot readily be applied to evolution in natural conditions, such as HIV-1 evolution within or between hosts. Current computational methods to infer fitness from population dynamics ignore linkage or suffer from serious computational costs (18–23).

Here we describe a method to infer selection from genetic time series data and demonstrate its ability to resolve linkage effects. We apply our method to reveal patterns of selection in intrahost HIV-1 evolution. Our approach is to efficiently quantify the probability of an evolutionary ‘path,’ defined by the set of all mutant allele frequencies at each time, using a path integral method derived from statistical physics (Methods). This expression can be analytically inverted to find the parameters that are most likely to have generated a path.

To define the path integral, we consider Wright-Fisher population dynamics with selection, mutation, and recombination, in the diffusion limit (24). Under an additive fitness model, the fitness of any individual is a sum of selection coefficients *s*_*i*_, which quantify the selective advantage of mutant allele *i* relative to wild-type. The probability of an evolutionary path is then a product of probabilities of changes in mutant allele frequencies at each locus between successive generations, including the influence of selection at linked loci.

Applying Bayes’ theorem leads to an analytical expression for the maximum *a posteriori* vector of selection coefficients ***ŝ*** corresponding to a path (Methods),

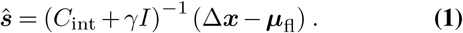

The integrated covariance matrix of mutant allele frequencies *C*_int_ accounts for the speed of evolution and linkage effects. Here *γ* quantifies the width of a Gaussian prior distribution for selection coefficients, and *I* is the identity matrix. Intuitively, the net change in mutant allele frequencies Δ***x*** and the integrated mutational flux ***µ***_fl_ determine whether the dynamics of a mutant allele appear to be beneficial or deleterious when linkage is ignored. Because equation (1) emerges from the likelihood of allele frequency trajectories, a subset of the full genotype distribution, we refer to it as the marginal path likelihood (MPL) estimate of the selection coefficients.

To test the ability of MPL to uncover selection, we analyzed data from simulations of a variety of evolutionary scenarios (Supplementary Information). Even in cases with strong linkage (Fig. 1A), MPL accurately recovers true selection coefficients (Fig. 1B). Further tests indicated that performance remains strong even when data is limited, an important practical consideration (Supplementary Fig. 1). Compared to existing methods of selection inference (18–23), MPL was the most accurate method in terms of both classification accuracy, measured by AUROC for classifying mutant alleles as beneficial or deleterious, and in the absolute error in inferred selection coefficients (Supplementary Fig. 2, Supplementary Information) across a range of simulated data sets. Due to the simplicity of equation (1), MPL was fastest among the methods that we compared, with a running time roughly 6 orders of magnitude faster than approaches that rely on iterative Monte Carlo methods.

**Fig. 1.**
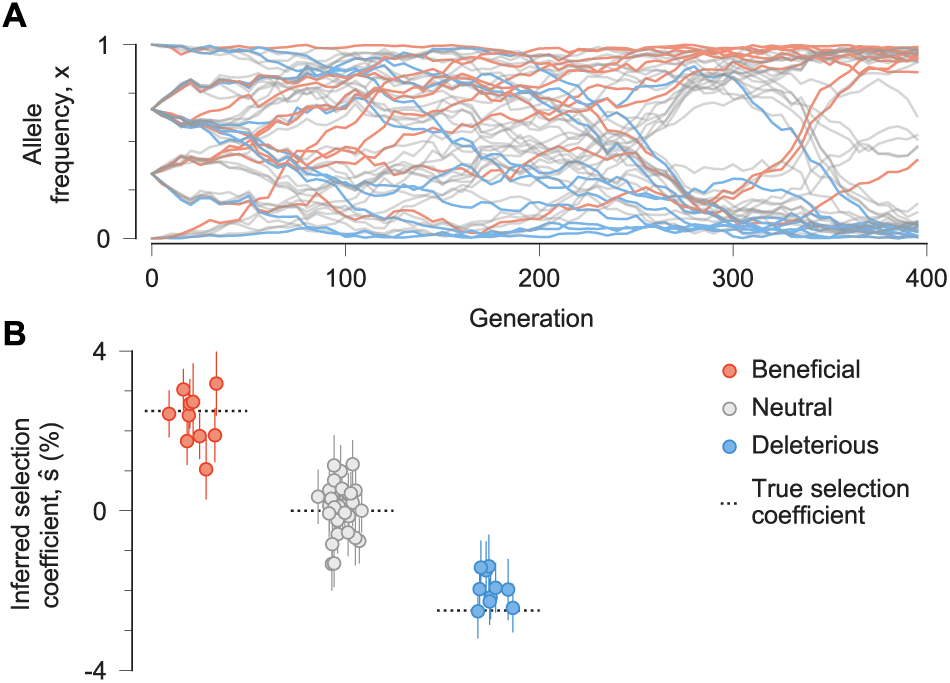
MPL accurately recovers selection from complex dynamics. **A**, Simulated allele frequency trajectories in a model with 10 beneficial, 30 neutral, and 10 deleterious mutant alleles. The initial population is a mix of three subpopulations with random mutations. Selection is challenging to discern from individual trajectories alone. **B**, Selection coefficients inferred by MPL are close to their true values. Error bars denote theoretical standard deviations for inferred coefficients. Simulation parameters are the same as those defined in Supplementary Fig. 1.

Next we applied MPL to study the intrahost evolution of HIV-1 and to resolve interactions between HIV-1 and the immune system. Identifying selective pressures on HIV-1 gives insight into the evolutionary dynamics leading to HIV-1 escape from immune control and the development of bnAbs, both of which are relevant for vaccine design.

We first examined longitudinal HIV-1 half-genome sequence data from 13 individuals where early-phase CD8^+^ T cell responses were comprehensively analyzed (25). In this group, 36.6% of the top 1% most beneficial mutations reported by MPL are nonsynonymous mutations in identified (25) CD8^+^ T cell epitopes (Fig. 2A). This is a 19-fold enrichment in mutations in T cell epitopes compared to expectations by chance (Methods). Reversions to clade consensus are also strongly beneficial. Nonsynonymous reversions outside of T cell epitopes are 14-fold enriched in this subset. Furthermore, nonsynonymous reversions within T cell epitopes are 326-fold enriched. These findings are compatible with past studies that have observed strong selection for T cell escape (8, 9, 26) and for reversions (9).

**Fig. 2.**
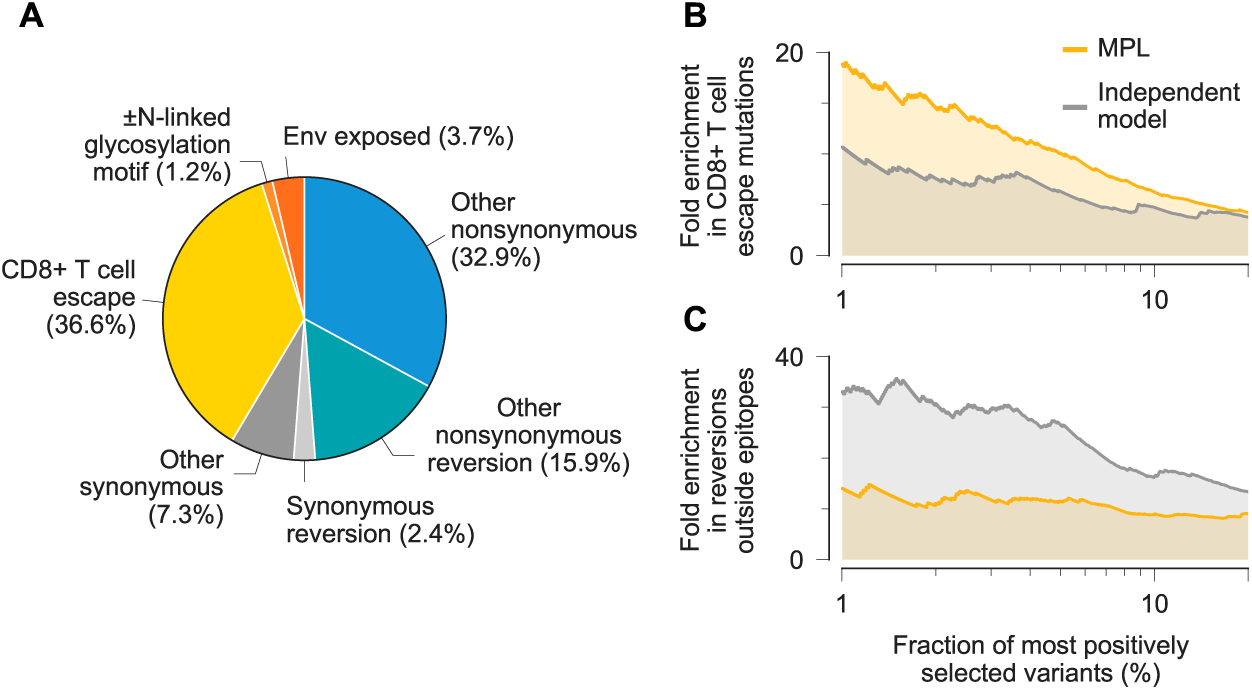
Patterns of strong selection in within-host HIV evolution. **A**, Among the top 1% most beneficial variants across individuals, mutations to escape from T cell-mediated immunity are especially common. **B**, Due to clonal interference between escape mutants, MPL identifies more escape variants to be strongly beneficial than an independent model which ignores covariance. **C**, In contrast, the independent model estimates an excess in the number of strongly beneficial reversions.

Resolving linkage leads to substantial differences in selection estimates. MPL places 1.76 times as many T cell escape mutations within the top 1% most beneficial mutations as an independent model that ignores linkage between mutant alleles (Fig. 2B). Conversely, MPL ranks 0.42 times as many nonsynonymous reversions outside of T cell epitopes to be strongly beneficial as the independent model does (Fig. 2C). These differences are explained by the collective resolution of genetic linkage effects, including clonal interference.

In order to dissect the contributions of linkage to estimates of fitness, we computed the pairwise effects Δ*ŝ*_*ij*_ of each variant *i* on the inferred selection coefficients for all other variants *j* (Methods). We defined Δ*ŝ*_*ij*_ as the difference between the estimated selection coefficient *ŝ*_*j*_ for variant *j* using all of the data and the value of *ŝ*_*j*_ when variant *i* is replaced by the transmitted/founder (TF) nucleotide at the same site, thereby removing the contribution to selection from linkage with variant *i*. Positive values of Δ*ŝ*_*ij*_ indicate that linkage with variant *i* increases the selection coefficient inferred for variant *j* (e.g., due to clonal interference between them). Negative values indicate that variant *i* decreases the selection coefficient inferred for variant *j* (e.g., due to hitchhiking).

Our analysis revealed that the vast majority of observed variants have essentially no effect on estimates of selection at other sites, but a small minority of highly influential variants have dramatic effects (Supplementary Figs. 3-4). Such highly influential variants are often ones that change rapidly in frequency, sweeping through the population and exerting strong effects on linked sites (Supplementary Fig. 5). Consistent with this observation, 40% of highly influential variants are putative CD8^+^ T cell escape mutations. Effects on estimated selection drop off sharply with increasing distance along the genome for most variants (Supplementary Fig. 6). However, the effects of highly influential variants routinely span across long genomic distances (Supplementary Fig. 4). Collectively, this data suggests that some highly influential variants are drivers of selective sweeps. Competition between such variants results in clonal interference.

A clear example of clonal interference is demonstrated in escape from a T cell response in individual CH77 targeting the Nef KF9 epitope (Fig. 3A). MPL infers strong positive selection for all escape variants. In contrast, when linkage is ignored escape variants that are lost are inferred to be neutral, and the magnitude of selection for 9040C is substantially decreased (Fig. 3B). Ignoring linkage thus leads to selection estimates that are qualitatively and quantitatively suspect. We observe similar instances of clonal interference in other epitopes (Supplementary Figs. 7-8). In the case of KF9, comfpetition between the different escape variants increases the estimated selection coefficient for each of them (Fig. 3C). Inrferred selection is also influenced by linkage with other mutations outside the KF9 epitope (Fig. 3C,D). For example, 9040C is inferred to be more beneficial due to its competition with the DI9 escape mutation 6021C. The selection coefficient for 9044G, in turn, is somewhat reduced due to positive linkage with 8719G, which is the dominant escape mutation in the nearby Env DR9 epitope.

**Fig. 3.**
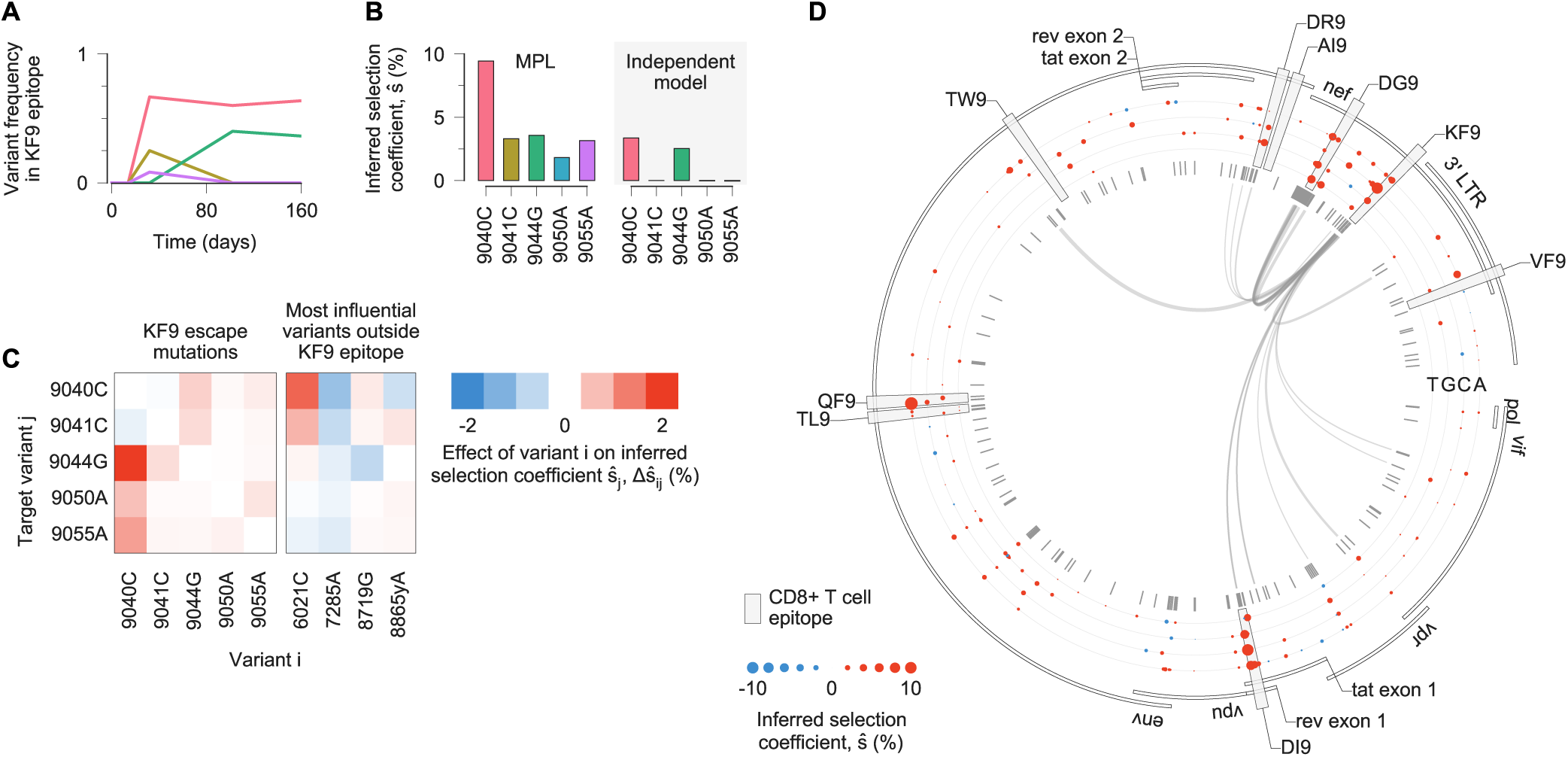
Estimates of selection coefficients for viral escape mutations must account for clonal interference. **A**, Multiple escape mutations appear in the viral population in the T cell epitope KF9, targeted by individual CH77, exhibiting clonal interference. **B**, Using the full half-genome-length sequence data as input, MPL infers that all KF9 escape variants are positively selected. In contrast, estimates based solely on the trajectories of individual variants only uncover substantial positive selection for the 9040C and 9044G variants that coexist at the final time point. Furthermore, the independent model estimates of selection are attenuated because of the failure to account for competition with other beneficial mutations, including other escape mutations within the same epitope. **C**, Linkage effects on inferred selection coefficients for KF9 escape mutations. Effects shown here are due to variants within the KF9 epitope and the top four most influential variants outside the KF9 epitope, defined as the variants *i* for which Σ_*j*_ |Δ*ŝ* _*ij*_ | is the largest. All of these influential variants lie within other T cell epitopes (6021C lies in DI9, 7285A in QF9, 8719G in DR9, and 8865yA in DG9). **D**, Inferred selection in the HIV-1 half-genome sequence for CH77. Inferred selection coefficients are plotted in tracks. Coefficients of TF nucleotides are normalized to zero. Tick marks denote polymorphic sites. Inner links, shown for sites connected to the KF9 epitope, have widths proportional to matrix elements of the inverse of the integrated covariance (see Eq. (1)).

Some strongly selected mutations lie in regions of Env that are exposed to antibodies, or in N-linked glycosylation motifs that affect the area of Env that is accessible to antibodies (Fig. 2A). However, these mutations are infrequent compared to others in T cell epitopes. One can also observe little positive selection in Env outside of T cell epitopes in the example of CH77 (Fig. 3D). Overall, selection for escape from antibody responses appears to be weaker or less frequent than CD8^+^ T cell-mediated selection.

We then asked whether strong antibody-mediated selection would be observed in individuals who generate bnAbs. To explore this question we studied HIV-1 evolution in individual CAP256, who eventually developed the VRC26 family of bnAbs (27, 28). This case is particularly challenging for inference because of a superinfection event 15 weeks after initial infection (Fig. 4A). This leads to strong and complex patterns of linkage as the superinfecting strain recombines and competes with the primary infecting strain (Fig. 4B, Supplementary Fig. 9). For this reason, ignoring linkage leads to poor selection inferences. Most (6 of 11) of the top 1% most beneficial mutations inferred by the independent model are from the background of the superinfecting strain and are synonymous. In contrast, none of the most beneficial mutations inferred by MPL are synonymous.

**Fig. 4.**
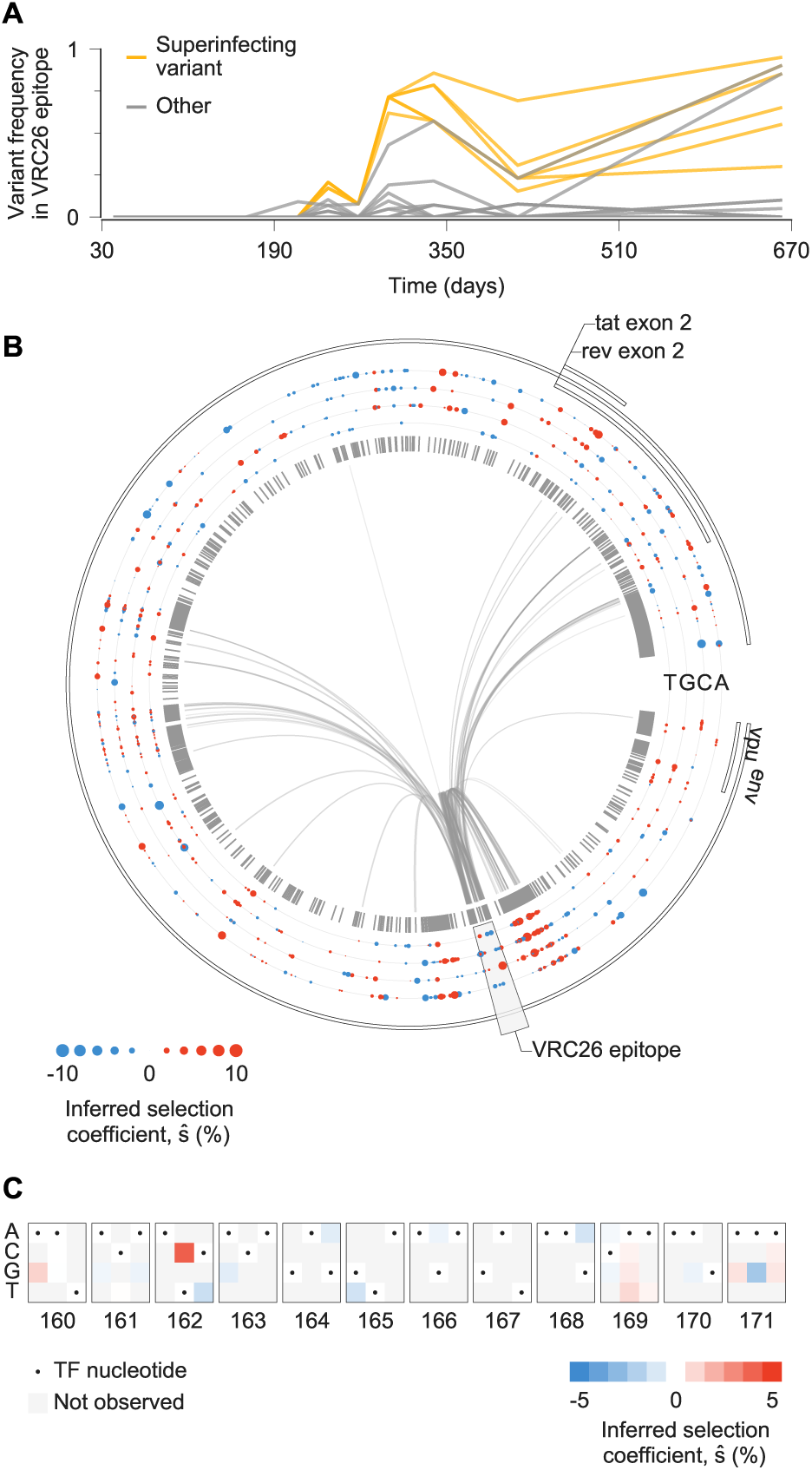
Complex patterns of selection in HIV-1 Env following superinfection in an individual who develops broadly neutralizing antibodies. **A**, Multiple variants, including several from the superinfecting strain of the virus, rise and fall in frequency within the epitope targeted by the VRC26 family of antibodies. **B**, Inferred selection in CAP256 HIV-1 half-genome sequences. Inferred selection coefficients are plotted in tracks. Coefficients of TF nucleotides are normalized to zero. Tick marks denote polymorphic sites. Inner links, shown for sites connected to the VRC26 epitope, have widths proportional to matrix elements of the inverse of the integrated covariance. Linkage is extensive due to the struggle for dominance in the viral population between the TF, superinfecting, and recombinant strains. **C**, Map of inferred selection within the VRC26 epitope, consisting of codons 160-171 in Env.

We found that selection for known VRC26 resistance mutations (27, 28) is modest (Fig. 4C). The most strongly selected mutation in the VRC26 epitope region is 6709C (*ŝ* = 0.041) in codon 162 in Env, a variant present in the superinfecting strain that completes an N-linked glycosylation motif which is absent from the primary infecting virus. However, this modification makes the virus more sensitive to VRC26 (27, 28). We observe selection against 6717T (*ŝ* = −0.012), corresponding to the Env 165L variant in the superinfecting strain. Reversion of this residue to V, the variant in the primary infecting strain, improves resistance to early VRC26 antibodies (28). We also observe modest positive selection for nonsynonymous variation at codon 169 in Env (maximum *ŝ*= 0.010), where mutations lead to complete resistance to VRC26 family antibodies (28). Thus, even the most strongly selected resistance mutations would fall outside of the top 5% most strongly selected mutations in the larger sample of 13 individuals.

Weak selection on the virus for antibody escape may in fact facilitate the development of bnAbs. Multiple escape variants, as well as variants that are sensitive to the antibody, can readily coexist for long times when escape is weakly selected. This coexistence increases the diversity of the viral population. Pressure on antibodies to bind to multiple variants can then select for breadth (29). Indeed, viral diversification has been observed to precede bnAb development (28, 30). In contrast, stronger pressure on the virus for escape could reduce viral diversity due to rapid fixation of beneficial escape variants and the elimination of sensitive ones.

Overall, our results reveal patterns of HIV-1 adaptation, including selection on the virus population accompanying the development of bnAbs, that were not possible to quantify using methods that cannot resolve genetic linkage. Furthermore, the scale of the data considered here is far beyond what existing methods that attempt to account for linkage can analyze. We anticipate that our method can also be widely applied to investigate selection in other evolving populations. Given the potential pitfalls of linkage-naive inference, our results call for a greater focus on resolving linkage effects in studies of selection.

## ACKNOWLEDGEMENTS

We thank A. K. Chakraborty, C. J. R. Illingworth, and J. G. Schraiber for helpful discussions and comments on the manuscript. The work of M.S.S., R.H.Y.L., and M.R.M. was supported by the Hong Kong Research Grants Council under grant number 16234716. R.H.Y.L. was also supported by Australia’s National Health and Medical Research Council under grant number APP1121643. J.P.B. acknowledges support from a Regents Faculty Fellowship provided by the UCR Academic Senate.

## AUTHOR CONTRIBUTIONS

All authors designed research, developed methods, analyzed data, interpreted results and wrote the paper.

## Methods

### Data and code availability

All data and computer code supporting the findings of this study are available on GitHub in the repository https://github.com/bartonlab/paper-MPL-inference.

### Evolutionary model

Our inference approach is based on the standard Wright-Fisher (WF) model of population genetics, which describes the stochastic dynamics of an evolving population of *N* individuals. Each individual is represented by a genetic sequence of length *L*. The population evolves in discrete, non-overlapping generations subject to the forces of selection, mutation, and recombination. For simplicity, we begin by describing the model with two alleles per locus, wild-type (WT) and mutant. Thus there are *M* = 2^*L*^ unique genotypes. Later we show that our approach readily generalizes to consider multiple alleles per locus.

The state of the population at a generation *t* is given by the genotype frequency vector ***z***(*t*) = (*z*_1_(*t*), *…*, *z*_*M*_ (*t*)), where *z*_*a*_(*t*) denotes the frequency of individuals with genotype *a*. Conditioned on ***z***(*t*), the probability that the genotype frequency vector in the next generation is ***z***(*t* + 1) is multinomial (31):

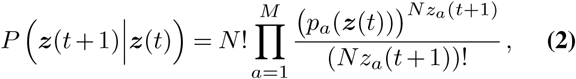

with

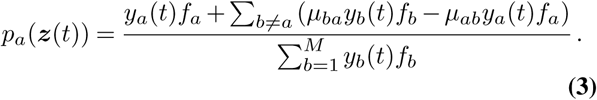

Here *f*_*a*_ denotes the fitness of genotype *a*, and *µ*_*ab*_ is the probability of genotype *a* mutating to genotype *b*. For simplicity we will assume at first that the mutation probability *µ* is the same at all loci, and that the probability of mutating from WT to mutant is the same as that from mutant to WT. In Eq. (3),

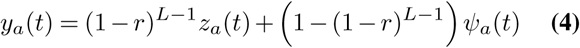

is the frequency of genotype *a* after recombination. Here *r* is the probability of recombination per locus per generation, and *Ψ*_*a*_(*t*) is the probability that randomly recombining any two individuals in the population results in an individual of genotype *a* (see Supplementary Information). Though Eq. (3) and Eq. (4) appear complex, they have an intuitive interpretation. The first term in Eq. (3) reflects the fact that fitter individuals reproduce more efficiently and are therefore more likely to be observed in future generations. Mutations, captured through the second term, lead to conversions from genotype *a* to other genotypes, and vice versa. The denominator in Eq. (3) provides an overall normalization and indicates that relative fitness is important: in order for a particular genotype to reliably grow in frequency, its fitness should be higher than the average fitness of all individuals in the population. The first term of Eq. (4) gives the proportion of individuals of genotype *a* not undergoing recombination, while the second term accounts for the net inward flow due to recombination from all other genotypes to genotype *a*.

For a population evolving under the WF model for *T* generations, the probability that the genotype frequency vector follows a particular evolutionary path (***z***(1), ***z***(2), *…*, ***z***(*T*)), conditioned on the initial state ***z***(0), is

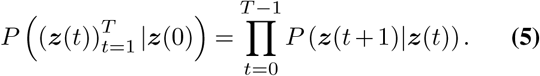

This expression is difficult to work with for parameter inference. This is due in part to the high dimensionality of the vector ***z***, which scales exponentially with the length of the genetic sequence. Thus, in the vast majority of real data sets, the sequence space is dramatically under-sampled. The functional form of Eq. (2) is also complex.

Our approach circumvents these issues by employing two approximations. First, we obtain a simplified version of Eq. (5) by using a path integral. Path integral expressions for evolutionary models have also been derived under different assumptions in past work (32–34), but they have not been widely applied for inference. We also assume that fitness is additive, such that the total fitness of each genotype *a* is just given by the sum of the selection coefficients *s*_*i*_ for mutant alleles at each locus *i*,

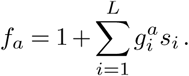

Here 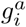 is 1 if genotype *a* has a mutant allele at locus *i* and 0 otherwise. These assumptions will substantially simplify the expression for Eq. (5).

### Path integral for mutant allele frequencies

In this section we will develop a simplified version of Eq. (5) defined at the level of allele frequencies rather than genotype frequencies. Later we will demonstrate that, if the fitness effects of mutations are additive as assumed above, this approach will lead to no loss of information for estimating the selection coefficients from data. We begin by using the WF dynamics above, which are defined for genotype frequencies, to compute the expected changes in frequency of mutant alleles. The mutant allele frequency *x*_*i*_ at locus *i* is

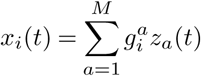

Following the assumptions above, and in the WF diffusion limit (31), one can show (Supplementary Information) that the probability density for mutant allele frequencies ***x***(*t*) = (*x*_1_(*t*), *x*_2_(*t*), *…*, *x*_*L*_(*t*)) follows a Fokker-Planck equation with a drift vector *d*_*i*_ and diffusion matrix *C*_*ij*_*/N* given by

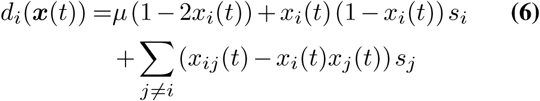

and

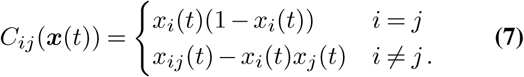

Here *x*_*ij*_ is the frequency of individuals in the population with mutant alleles at both loci *i* and *j*. The drift vector describes the expected change in mutant allele frequencies in time. Note that the last term in Eq. (6) quantifies linked selection, i.e., how the dynamics of mutant allele frequencies are affected by the average genetic background on which they appear. The drift vector should not be confused with *genetic drift*, the fluctuation in allele frequencies due to the inherent stochasticity of replication, which is instead described by the diffusion matrix. The diffusion matrix is simply the co-variance matrix of mutant allele frequencies divided by the population size *N*. It therefore depends on the double mutant frequencies *x*_*ij*_, but we will use the shortened notation *C*_*ij*_(***x***(*t*)) for brevity.

Applying standard methods from statistical physics (35), the Fokker-Planck equation can be converted into a path integral that quantifies the probability density for ‘paths’ of mutant allele frequencies (***x***(1), ***x***(2), *…*, ***x***(*T*)). This expression will allow us to efficiently estimate the parameters that are most likely to have generated a specific path (see Supplementary Information for details). The probability for a path is

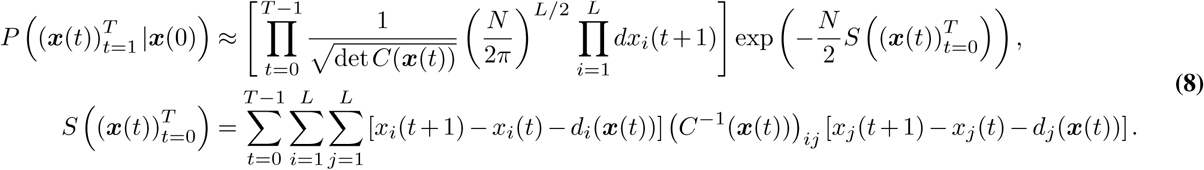

In the language of physics, 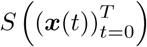 is referred to as the *action*. The population size *N* is analogous to the inverse temperature in statistical physics. The action penalizes deviation of the change in mutant frequencies between generations from the expectation given by the drift vector at the previous generation. This is normalized by the diffusion matrix, which quantifies the magnitude of typical changes in mutant frequencies due to random replication alone (i.e., genetic drift). The path integral equation in Eq. (8) follows the Itô convention.

### Marginal path likelihood (MPL) estimate of the selection coefficients

Given an observed path of mutant allele frequencies, we can employ Bayesian inference to determine the maximum *a posteriori* selection coefficients ***ŝ*** corresponding to the data, assuming that the population size *N* and mutation probability *µ* are known. In practice, our data consists of sets of genetic sequences obtained from a population at multiple time points *t*_*k*_, *k* ∈ {0, 1, *…*, *K*}. These sequences can be used to compute the path of mutant allele frequencies (***x***(*t*_0_), ***x***(*t*_1_), *…*, ***x***(*t*_*K*_)) as well as the double mutant frequencies *x*_*ij*_(*t*_*k*_), which also appear in Eq. (8). We will assume that the observed mutant allele frequencies are equal to the true ones, which simplifies the inference procedure. Our tests indicate that our results are robust to errors in the frequencies due to finite sampling (see Supplementary Fig. 1). Future work will relax this assumption.

In total, the posterior probability of the selection coefficients ***s*** = (*s*_1_, *s*_2_, *…*, *s*_*L*_) given the observed path (***x***(*t*_0_), ***x***(*t*_1_), *…*, ***x***(*t*_*K*_)) is

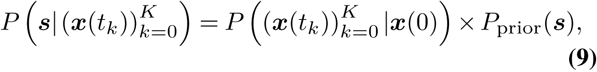

where 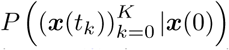 is the probability of the path (given by Eq. (8), but extended to arbitrary sampling times as shown in Supplementary Information) and *P*_prior_(***s***) is the prior probability for the selection coefficients. Eq. (9) is a complicated expression of the mutant allele frequencies. However, solving for the selection coefficients that maximize the posterior probability is straightforward because Eq. (8) is a Gaussian function of the selection coefficients. Taking *P*_prior_(***s***) to be a Gaussian distribution with mean zero and covariance matrix *σ*^2^*I*, where *I* is the identity matrix, the selection coefficients that maximize Eq. (9) are given by

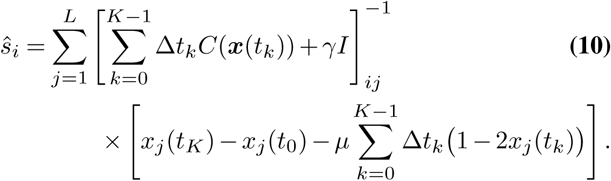

Here *γ* = 1*/Nσ*^2^, and Δ*t*_*k*_ = *t*_*k*+1_ − *t*_*k*_. We refer to Eq. (10) as the *MPL estimate* of selection coefficients. Because of the Gaussian form of Eq. (9), the maximum *a posteriori* estimates of the selection coefficients are the same as their pos-terior means.

Eq. (10) can be readily interpreted. Let us start by considering the vector term in the “numerator” of Eq. (10) that multiplies the matrix inverse. Here the first terms quantify how the frequency of each mutant allele has changed between the initial and final generations. Naturally, alleles that increase in frequency over time are more likely to be beneficial. The remaining terms quantify the integrated mutational flux, i.e., population flow from mutant to WT (or vice versa) due to mutation. Net mutational flux from mutant to WT is also associated with higher fitness for the mutant allele. This is because this indicates that the mutant state maintained higher frequency than the WT over the trajectory, despite the force of mutation that drives the frequencies toward the same value. Together, these terms in the numerator of Eq. (10) determine whether a mutant allele is inferred to be beneficial or deleterious, at least when the off-diagonal elements of the matrix that it multiplies are zero.

While the numerator of Eq. (10) roughly determines the sign of selection, the denominator determines the strength of the inferred selection coefficient.Let us refer to 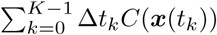 as the integrated covariance matrix. From Eq. (7) we see that the entries of *C*_*ij*_(***x***(*t*)) are small when the mutant frequency is near the boundaries (0 or 1). Thus, the dominant contribution to the integrated covariance matrix comes from points on the path where the mutant frequency is far from the boundaries. If selection is strong, so that the mutant allele is much fitter than the WT (or vice versa), then we expect that a large portion of the path will be spent with the mutant allele frequency close to the boundary. In such cases the diagonal part of the integrated covariance will be small, and we correctly infer strong selection. The prior distribution for the selection coefficients simply adds a constant to the diagonal of the integrated covariance, which both shrinks selection estimates toward zero and ensures that the matrix is invertible. The off-diagonal terms of the integrated covariance matrix capture linkage effects, that is, how much of the change in the mutant frequency at a locus can be attributed to the average sequence background on which the mutant appears.

### Equivalence of genotype- and allele-level analyses

In the preceding section we derived an estimate for the selection coefficients most likely to have generated an observed evolutionary path. To do this we used an expression for the likelihood of a path of mutant allele frequencies that depended on the mutant frequencies *x*_*i*_(*t*) and their pairwise correlations *x*_*ij*_(*t*), but not on higher order correlations of the full genotype distribution. However, the WF dynamics is defined at the level of genotypes.

It can be shown that the use of Eq. (8) does not result in any loss in information beyond the approximations inherent in the WF diffusion limit. In the WF diffusion limit, the same steps as those applied to derive Eq. (8) can be performed for the genotype frequencies (see Supplementary Information). This results in a path integral expression that quantifies the probability density of genotype frequency paths. As in the allele-level analysis, the estimated selection coefficients are those that maximize

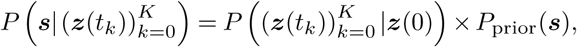

where 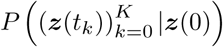 is the probability density of the genotype frequency path. The full expression is more complicated, and less transparent, than the allele-level equivalent. Nonetheless, one can show that the expression for the selection coefficients that maximize the posterior probability above is exactly the same as Eq. (10). Full details of this derivation are given in the Supplementary Information. This result is important because it shows that, following the assumptions of the WF diffusion limit and assuming that the fitness effects of mutations are additive, higher order mutational correlations contain *no further information* about the fitness effects of mutations.

### Extension to multiple alleles per locus and asymmetric mutation probabilities

The MPL framework extends readily to models with ℓ alleles per locus, as well as asymmetric mutation probabilities. Let *x*_*i,α*_(*t*) denote the frequency of allele *α* at locus *i* at generation *t*, and denote *µ*_*αβ*_ as the mutation probability per locus from allele *α* to allele *β*. Now, the trajectory of allele frequency vectors is (***x***(*t*_0_), ***x***(*t*_1_), *…*, ***x***(*t*_*K*_)), where ***x***(*t*_*k*_) = (*x*_1,1_(*t*_*k*_), *x*_1,2_(*t*_*k*_), *…*, *x*_1,*ℓ*_(*t*_*k*_), *x*_2,1_(*t*_*k*_), *…*, *x*_*L,ℓ*_(*t*_*k*_)). Following parallel arguments to before (see Supplementary Information), the MPL estimate of the selection coefficient *ŝ*_*i,α*_ for allele *α* at locus *i* is

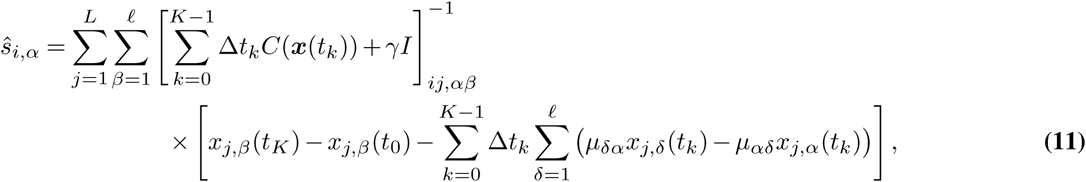

where *γ* = 1*/Nσ*^2^ as before. Off-diagonal entries of the covariance matrix *C*(***x***(*t*_*k*_)) are given by

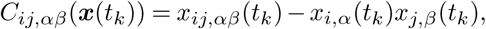

where *x*_*ij,αβ*_(*t*_*k*_) is the frequency of sequences with alleles *α* and *β* at loci *i* and *j*, respectively, at time *t*_*k*_.

### Data and code

Raw data and code used in our analysis is available in the GitHub repository located at https://github.com/bartonlab/paper-MPL-inference. This repository also contains Jupyter notebooks that can be run to reproduce the results presented here.

### Simulation data

We implemented the Wright-Fisher model with discrete generations in Python. We used this program to record 100 evolutionary histories each for three different choices of the underlying parameters. Parameter values are detailed in Supplementary Figs. 1 and 2. For all simulations we assumed only two alleles per site. The simulation code, code for analysis, and original simulation data are contained in the GitHub repository.

### Other time-series inference methods

The independent model that we compared MPL against in the main text is a single locus (SL) variant of MPL in which the off-diagonal elements of the integrated covariance matrix are set to zero.

The seven additional time series-based inference methods that we compared MPL against are FIT (36), LLS (37), CLEAR (38), EandR (39), ApproxWF (40), WFABC (41), and IM (42). Where available, we used the scripts provided by the authors. Some of these methods required preprocessing of the time-series data to obtain valid estimates of selection coefficients. See Supplementary Information for full details on implementation and data processing.

### Patient cohort

We studied HIV-1 sequence data obtained from a 14 individuals recruited under the CHAVI 001 and CAPRISA 002 studies in the United States, Malawi, and South Africa. The locations of CD8^+^ T cell epitopes were experimentally (43) or computationally (44) determined in 13 of the 14 individuals. In the remaining individual, CAP256, experimental studies identified the VRC26 family of antibodies and mapped the epitope location on Env (45).

### HIV-1 sequence data

Multiple sequence alignments of HIV-1 nucleotide sequences for all individuals were obtained from the Los Alamos National Laboratory HIV Sequence Database (www.hiv.lanl.gov; accessed October 19, 2018). Sequences labeled as problematic were not downloaded. For the 13 individuals with identified T cell epitopes, sets of 3^*1*^ and 5^*1*^ half-genome sequences were obtained, which were approximately 4500 bp in length. Only Env sequences were available for CAP256 (approximately 2500 bp in length). All sequences were aligned with the HXB2 reference sequence (GenBank accession number K03455) for numbering, and with clade consensus sequences to determine reversions, using the Los Alamos National Laboratory HIVAlign tool (46).

In order to assure sequence quality, we 1) removed sequences with ≥200 gaps with respect to clade consensus, 2) removed sites with >95% gaps in the alignment, 3) imputed ambiguous nucleotides, and 4) removed time points where <4 sequences were sampled, or where the time gap between successive samples exceeded 300 days. During this process we also determined the number of reading frames in which each substitution was nonsynonymous, whether it occurred within an identified CD8^+^ T cell epitope that was actively targeted during the time for which sequence samples were available, whether it occurred within the exposed surface of Env (using surface residues as identified in ref. (47)), and whether it may have plausibly affected Env glycosylation by completing or disrupting an N-linked glycosylation motif. These analyses were performed using custom Python scripts available in the GitHub repository.

Variant indices were labeled relative to the standard HXB2 reference sequence of HIV-1. Insertions relative to HXB2 are labeled with lowercase alphabetical indices per standard conventions (48). For example, if three nucleotides were inserted relative to HXB2 after site 1, these would be labeled 1a, 1b, and 1c, respectively.

### Enrichment analyses

We used fold enrichment values to determine the relative excess or lack of particular types of mutations among the HIV-1 variants that were inferred to be the most beneficial. For a given set of *N*_sel_ selected mutations (e.g., corresponding to the top 1% most beneficial), we computed the number *n*_sel_ of these mutations that have a particular property. This may represent, for example, the number of nonsynonymous mutations within identified CD8^+^ T cell epitopes, or the number of nonsynonymous reversions. The ratio *n*_sel_*/N*_sel_ then represents the fraction of the selected mutations having the specified property. This number was compared with *n*_null_*/N*_null_, where *N*_null_ is the total number of non-TF variants across all individuals and sequencing regions of the HIV-1 genome, and *n*_null_ is the number of these variants that share the specified property.

The fold enrichment of the selected set for a specified property is then naturally defined as (*n*_sel_*/N*_sel_)*/*(*n*_null_*/N*_null_). A fold enrichment value greater than one indicates a larger proportion of mutants in the selected set that have the given property than expected by chance, while a value less than one indicates a smaller proportion than expected by chance.

### Selection inference with MPL

We implemented the MPL method as described above in C++ and applied it to infer selection coefficients from the HIV-1 sequence data and from simulations. The original code used for inference is included in the GitHub repository. For the HIV-1 data, we assumed a regularization strength of *γ* = 5.

We also used mutation probabilities estimated in (49) as input. Mutation probabilities to and from gap states, representing deletions and insertions, respectively, were assumed to be very small (*µ* = 10^−9^). For the simulated data, we used a smaller regularization strength of *γ* = 1 due to the greater sampling depth. In order to increase robustness, we assumed that the true underlying allele frequency trajectories were piecewise linear and replaced the sums over time in Eq. (11) with integrals. Following the assumption of piece-wise linearity, these integrals can be computed analytically. Specifically, the contribution of the mutational term to the numerator is then

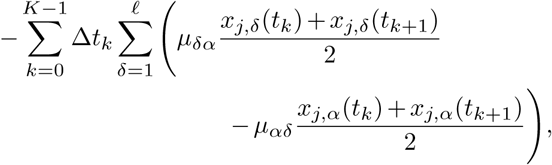

the diagonal terms of the integrated covariance matrix are

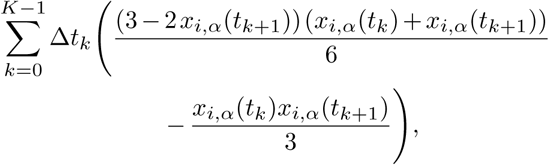

and the off-diagonal terms of the integrated covariance matrix are

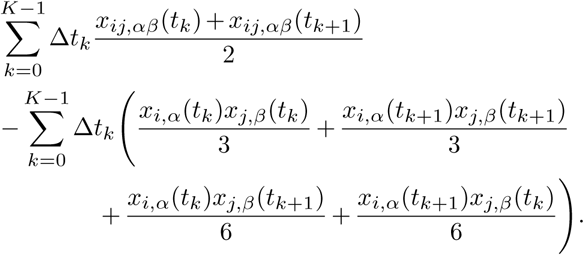

After selection coefficients were inferred, we normalized them such that the transmitted/founder (HIV-1) or wild-type (simulation) allele had a selection coefficient of zero.

### Calculation of effects of linkage on inferred selection

Due to the inverse of the integrated covariance matrix in Eq. (10), the selection coefficients estimated by MPL are affected by linkage. In order to quantify the effects of linkage on inferred selection during HIV-1 evolution, we computed the pairwise effects Δ*ŝ*_*ij*_ of each variant *i* on selection for other variants *j*, as described in the main text. Here, for ease of notation, each effective index *i* or *j* represents a single non-TF nucleotide at a particular site on the genome. That is, the indices incorporate both the label for the locus and for the allele.

To compute Δ*ŝ*_*ij*_, we iteratively select each nucleotide at each site, which together are represented by the index *i*, and generate a modified version of the sequence data in which variant *i* is replaced by the TF nucleotide at the same site. In this way, linkage between the masked variant *i* and all other variants *j* is eliminated. We then infer the selection coefficients again for all variants *j* using the data where variant *i* has been replaced by TF, denoted as 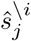. Then we define

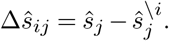

Positive values of Δ*ŝ*_*ij*_ thus indicate that linkage with variant *i* increases the selection coefficient inferred for variant *j*. This may be due, for example, to clonal interference between variants *i* and *j*. Negative values indicate that variant *i* decreases the selection coefficient inferred for variant *j*. This may occur, for example, if variant *j* hitchhikes on a beneficial genetic background that includes variant *i*.

---

**Supplementary Fig. 1.**
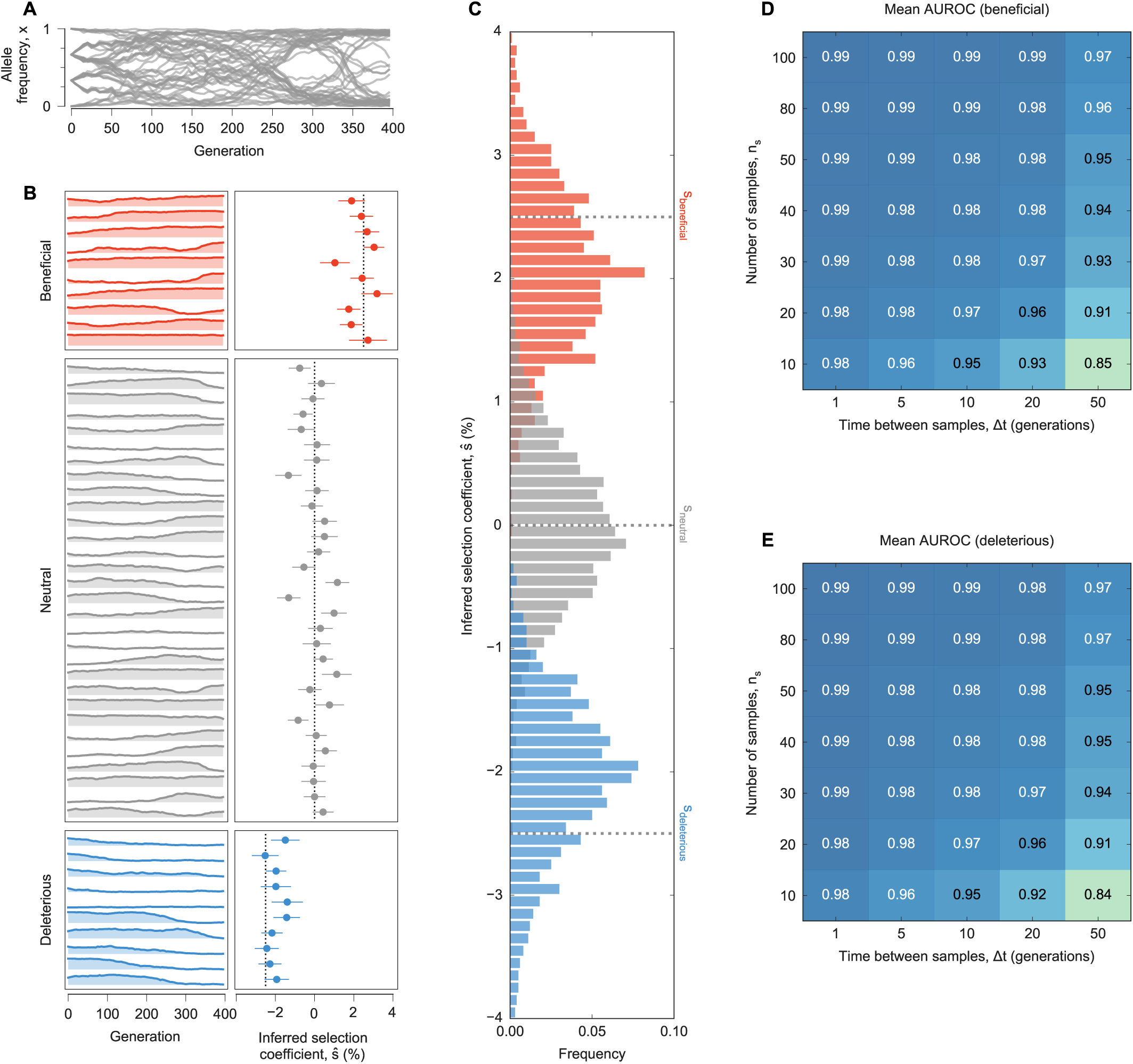
MPL accurately recovers selection coefficients from complex simulated evolutionary trajectories. **A**, Trajectories of mutant allele frequencies over time exhibit complex dynamics in a WF simulation with a simple fitness landscape. **B**, Separate views of individual trajectories for beneficial, neutral, and deleterious mutants (*left panel*) and inferred selection coefficients (*right panel*). Note that many neutral mutations exhibit temporal variation similar to beneficial or deleterious mutations. MPL estimates the underlying selection coefficients used to generate these trajectories and distinguishes between beneficial, neutral, and deleterious mutations, using equation (10). Dashed lines mark the true selection coefficients. **C**, Distributions of selection coefficient estimates across 100 replicate simulations with identical parameters in the special case of perfect sampling. MPL is also robust to finite sampling constraints, accurately classifying beneficial (**D**) and deleterious (**E**) mutants even when the number of sequences sampled per time point *n*_*s*_ is low, and the spacing between time samples Δ*t* is large. *Simulation parameters. L* = 50 loci with two alleles at each locus (mutant and WT): 10 beneficial mutants with *s* = 0.025, 30 neutral mutants with *s* = 0, and 10 deleterious mutants with *s* = − 0.025. Mutation probability *µ* = 10^−3^, population size *N* = 10^3^. Initial population composed of approximately equal numbers of three random founder sequences, evolved over *T* = 400 generations.

**Supplementary Fig. 2.**
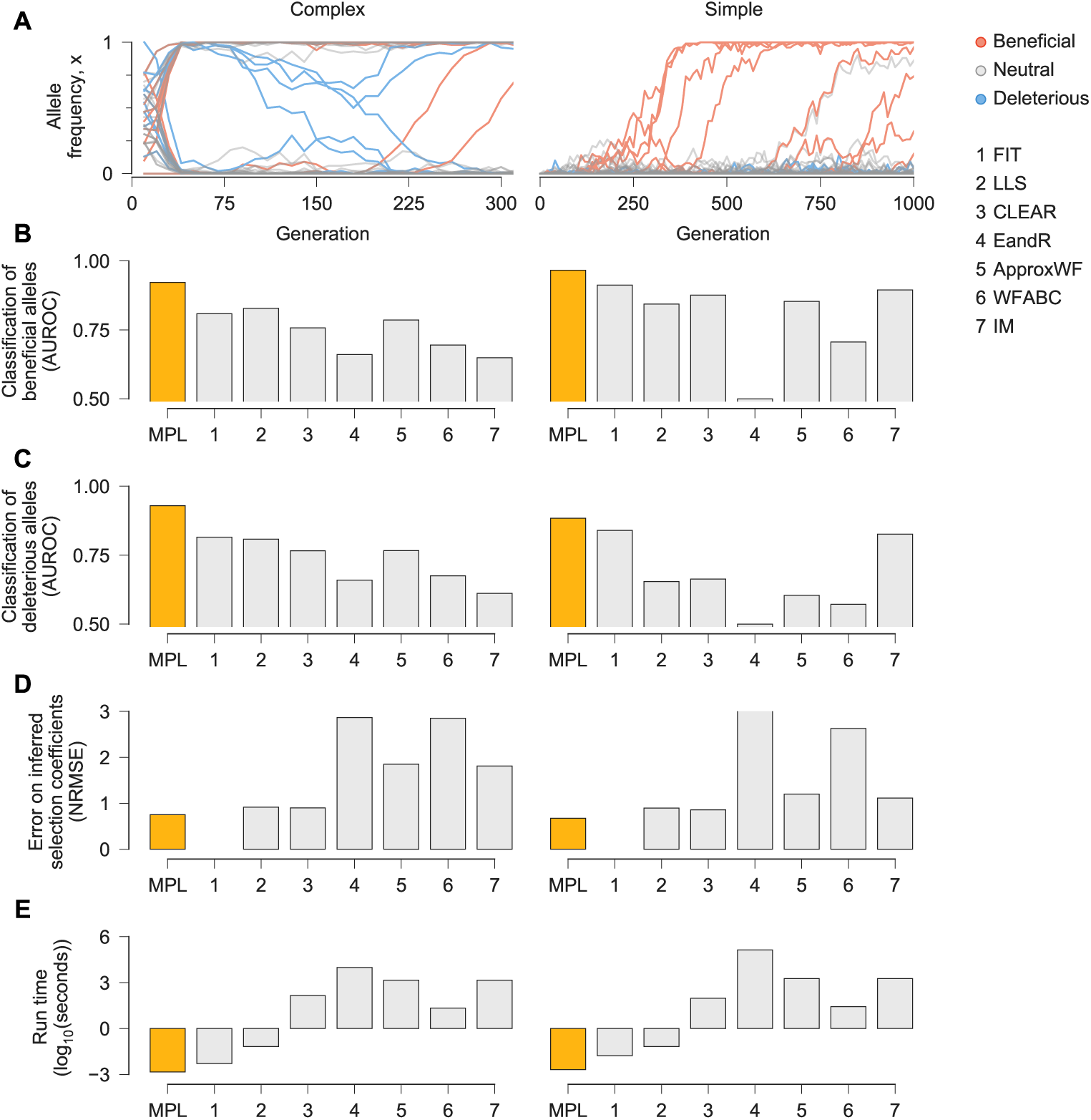
MPL compares favorably with state-of-the-art methods. We compared the ability of MPL and existing methods to infer selection from simulated test data that was rich with interference patterns and linkage, as shown in representative allele frequency trajectories (**A**). In order to evaluate robustness to finitely sampled data, we selected *n*_*s*_ = 100 sequences per time point for inference, with sampling time points separated by Δ*t* = 10 generations. Performance was evaluated by comparing the successful classification of beneficial (**B**) and deleterious (**C**) mutations, error in the estimated selection coefficients (**D**), and run time (**E**), averaged over 100 replicate simulations with identical parameters. MPL achieves the highest performance in terms of classification and estimation accuracy, and in run time. Note that FIT does not specifically estimate selection coefficients. *Simulation parameters. L* = 50 loci with two alleles at each locus (mutant and WT): 10 beneficial mutants (*s* = 0.1 for Complex, *s* = 0.025 for Simple), 30 neutral mutants (*s* = 0 for both scenarios), and 10 deleterious mutants (*s* = − 0.1 for Complex, *s* = − 0.025 for Simple). Mutation probability *µ* = 10^−4^, population size *N* = 10^3^. For the Complex case, the initial population is composed of equal numbers of five random founder sequences, evolved over *T* = 310 generations. Recorded trajectory used for inference begins at generation 10. For the Simple case, the initial population begins with all WT sequences, evolved over *T* = 1000 generations.

**Supplementary Fig. 3.**
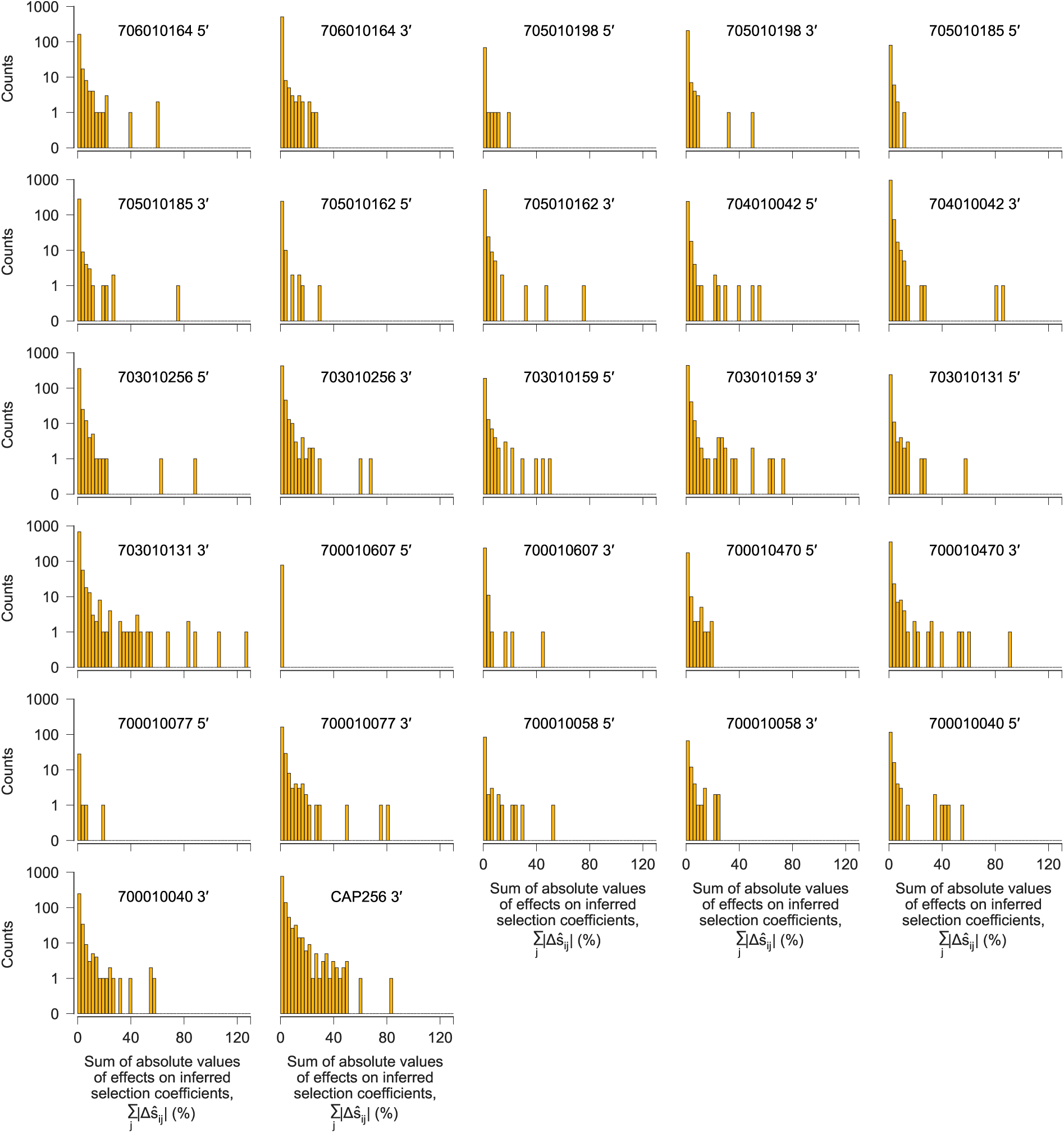
Most genetic variants have little effect on inferred selection at other sites, but a small minority have strong effects. After computing the pairwise effects Δ*ŝ*_*ij*_ of each variant *i* on the inferred selection coefficient for each other variant *j*, referred to as the target, we summed the absolute value of the Δ*ŝ*_*ij*_ values over all target variants *j* to quantify the influence of each variant *i* on selection at other sites. One histogram is shown for each sequencing region, for each individual. For the vast majority of variants, the total effect on selection at other sites is near zero. However, a small minority have strong effects. We defined a variant to be ‘highly influential’ if the sum of the absolute values of the Δ*ŝ*_*ij*_ over all targets *j* was larger than 0.4 (= 40%).

**Supplementary Fig. 4.**
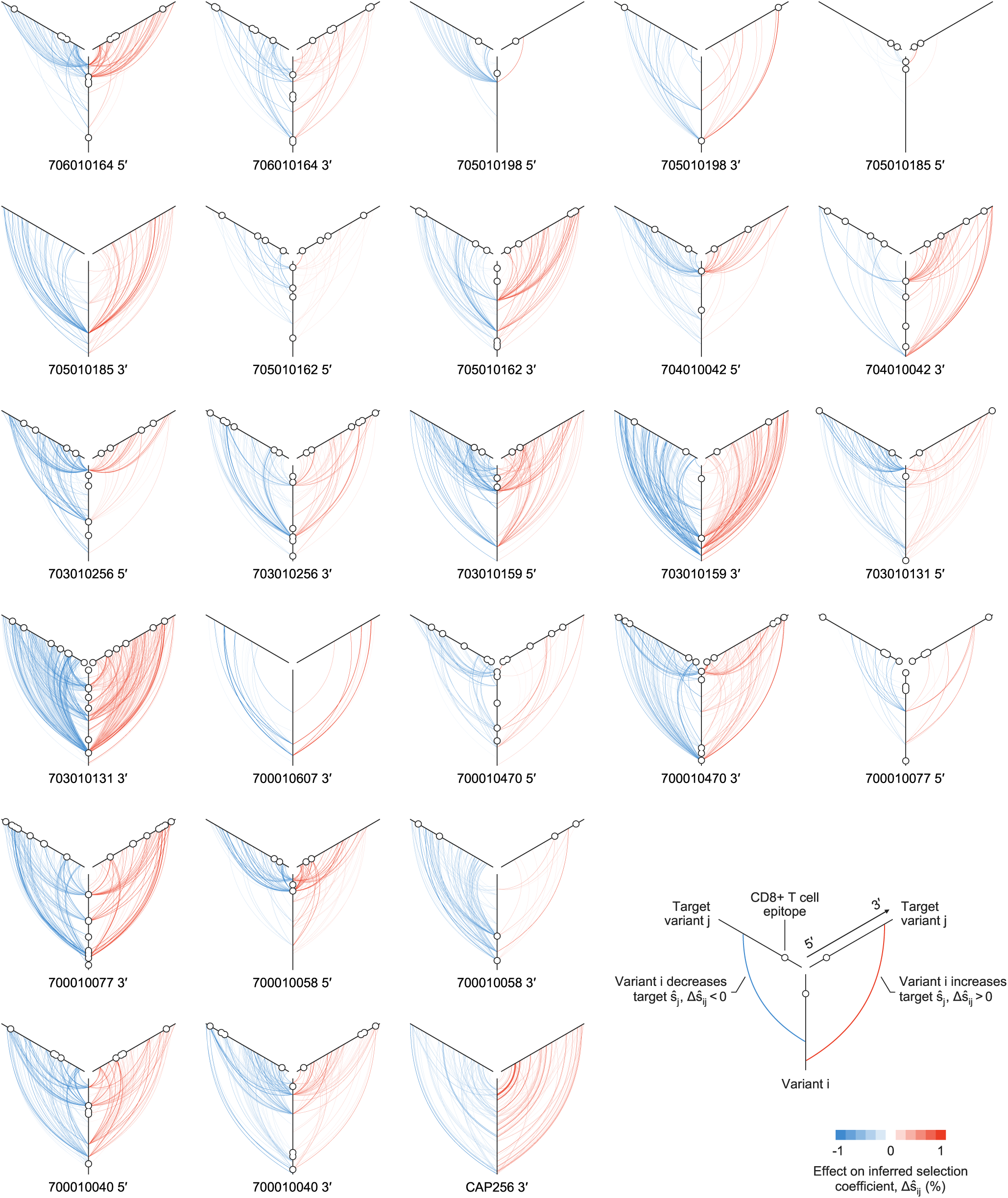
Variants that strongly influence inferred selection at other sites often act across large genomic distances. Plot of all linkage effects on inferred selection coefficients Δ*ŝ*_*ij*_ for which |Δ*ŝ*_*ij*_ | >0.004. One plot is shown for each sequencing region, for each individual. These strong effects of linkage on inferred selection coefficients can act at long range across the genome. Approximately 40% of highly influential variants, characterized by strong effects on inferred selection at other sites, lie within identified CD8^+^ T cell epitopes. The 5′ region for individual 700010607 is not shown because no Δ*ŝ*_*ij*_ values are larger than the cutoff.

**Supplementary Fig. 5.**
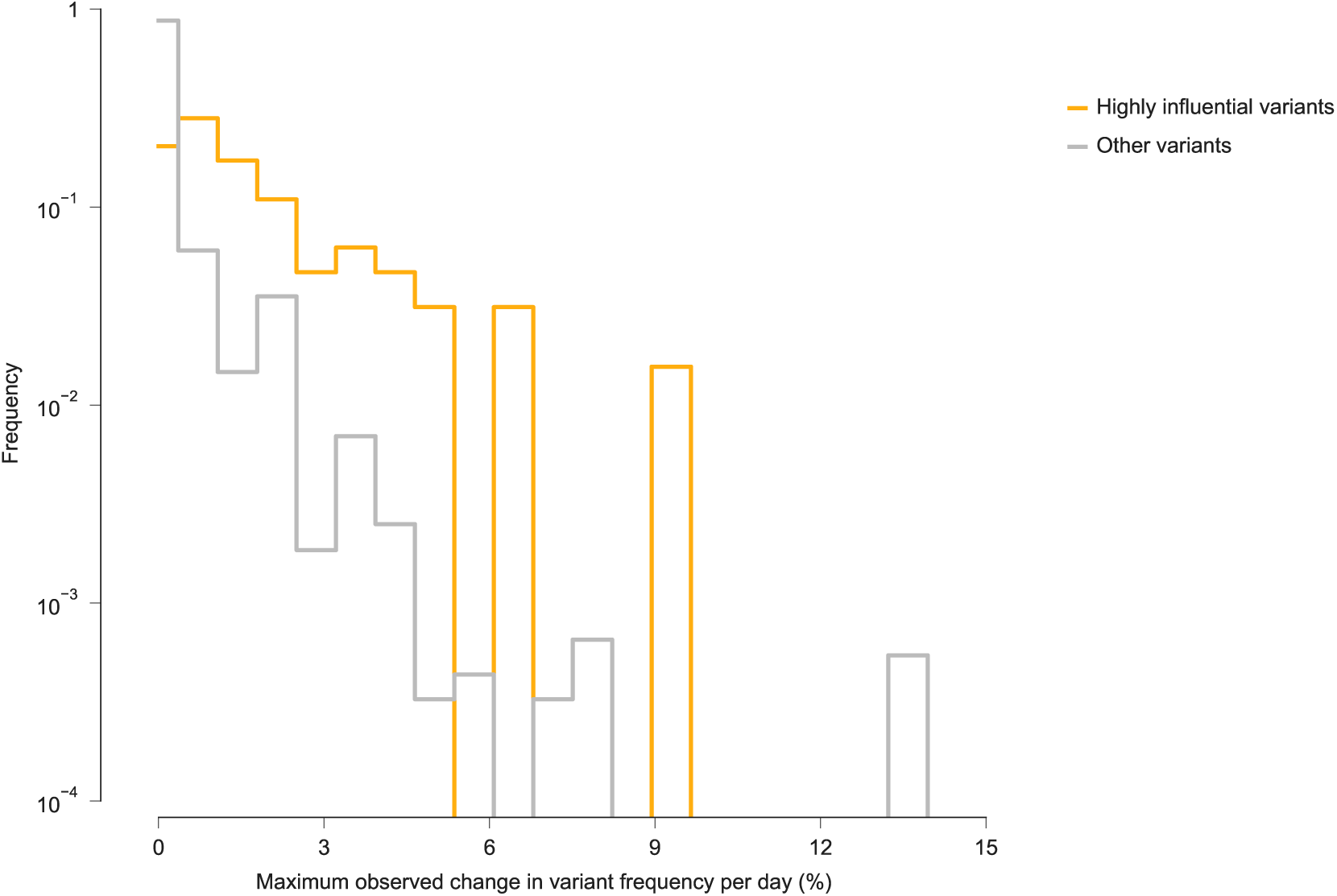
Highly influential variants are far more likely than other variants to change rapidly in frequency. For all genetic variants across all individuals and genomic regions considered in this study, we computed the maximum change in frequency per day between successive sequence samples. For most variants (those for which Σ_*j*_ | Δ *ŝ*_*ij*_| ≤ 0.4), the maximum change in frequency per day is less than 1%. Highly influential variants, which have large effects on inferred selection coefficients at other sites Σ_*j*_ | Δ *ŝ*_*ij*_| > 0.4), are much more likely than other variants to change rapidly in frequency.

**Supplementary Fig. 6.**
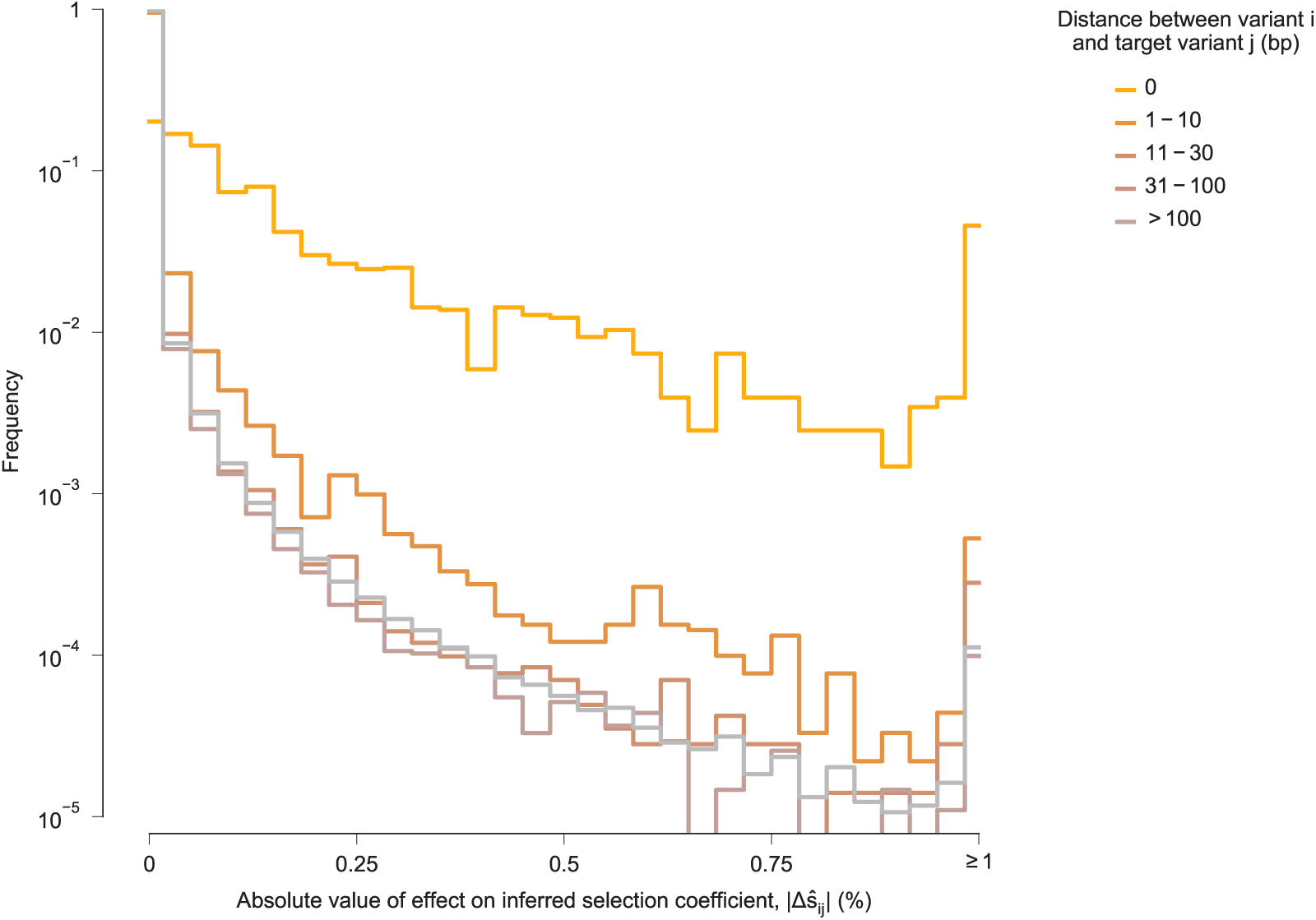
For most variants, effects on inferred selection coefficients for other variants are stronger at smaller genomic distances. Histogram of the absolute value of linkage effects on inferred selection coefficients for other variants |Δ*ŝ* _*ij*_ |, divided into subgroups based on the distance along the genome between variant *i* and target variant *j*. Consistent with intuition, the large effects on inferred selection coefficients occur most frequently for different variants that occur at the same site on the genome (i.e., distance equal to zero). “Interactions” between such variants are necessarily perfectly competitive, because only a single nucleotide is allowed at each position in the genetic sequence. For most variants, stronger linkage effects on inferred selection coefficients are more frequently observed for other variants within a distance of 10 base pairs (bp). Large linkage effects for pairs of variants within a distance of 30 bp, the approximate length of a linear T cell epitope, occur appreciably more frequently than for pairs of variants at greater genomic distances. However, there is little difference in the distribution of linkage effect sizes for pairs of variants that are between 31 bp and 100 bp apart compared to pairs of variants that are more than 100 bp apart. Nonetheless, some strong linkage effects on inferred selection are observed at long genomic distances (see Supplementary Fig. 4).

**Supplementary Fig. 7.**
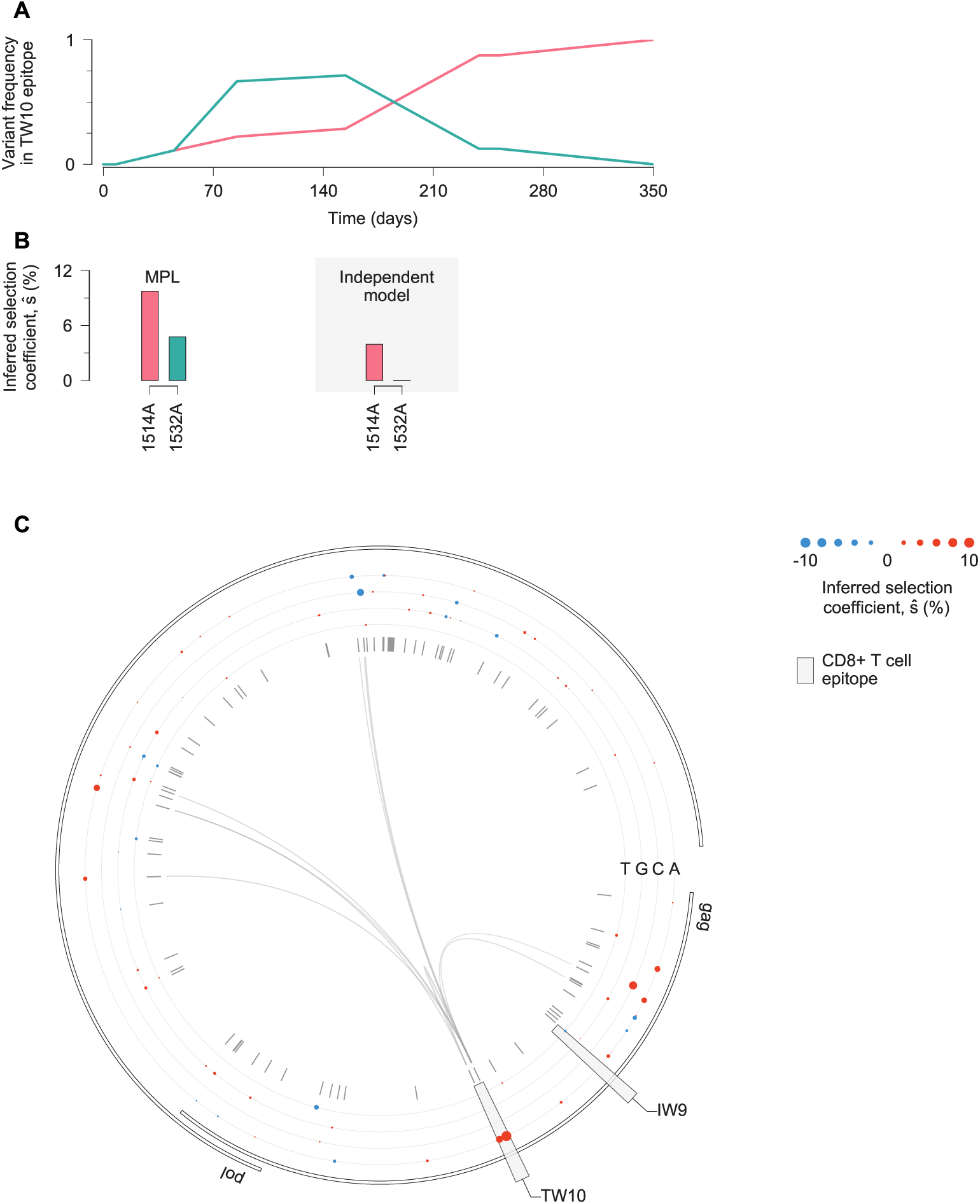
Estimates of selection coefficients in a simple example of clonal interference. **A**, Two escape mutations arise in the TW10 epitope targeted by individual CH58 and compete for dominance. **B**, MPL infers that both TW10 escape variants are positively selected. Estimates based on trajectories of individual variants only infer substantial positive selection for the 1514A variant that fixes. The magnitude of selection inferred with the independent model is also smaller than that inferred by MPL. **C**, Inferred selection in the HIV-1 5′ half-genome sequence for CH58. Inferred selection coefficients are plotted in tracks. Coefficients of transmitted/founder nucleotides are normalized to zero. Tick marks denote polymorphic sites. Inner links, shown for sites connected to the TW10 epitope, have widths proportional to matrix elements of the inverse of the integrated covariance. Linked sites affect selection estimates within the epitope.

**Supplementary Fig. 8.**
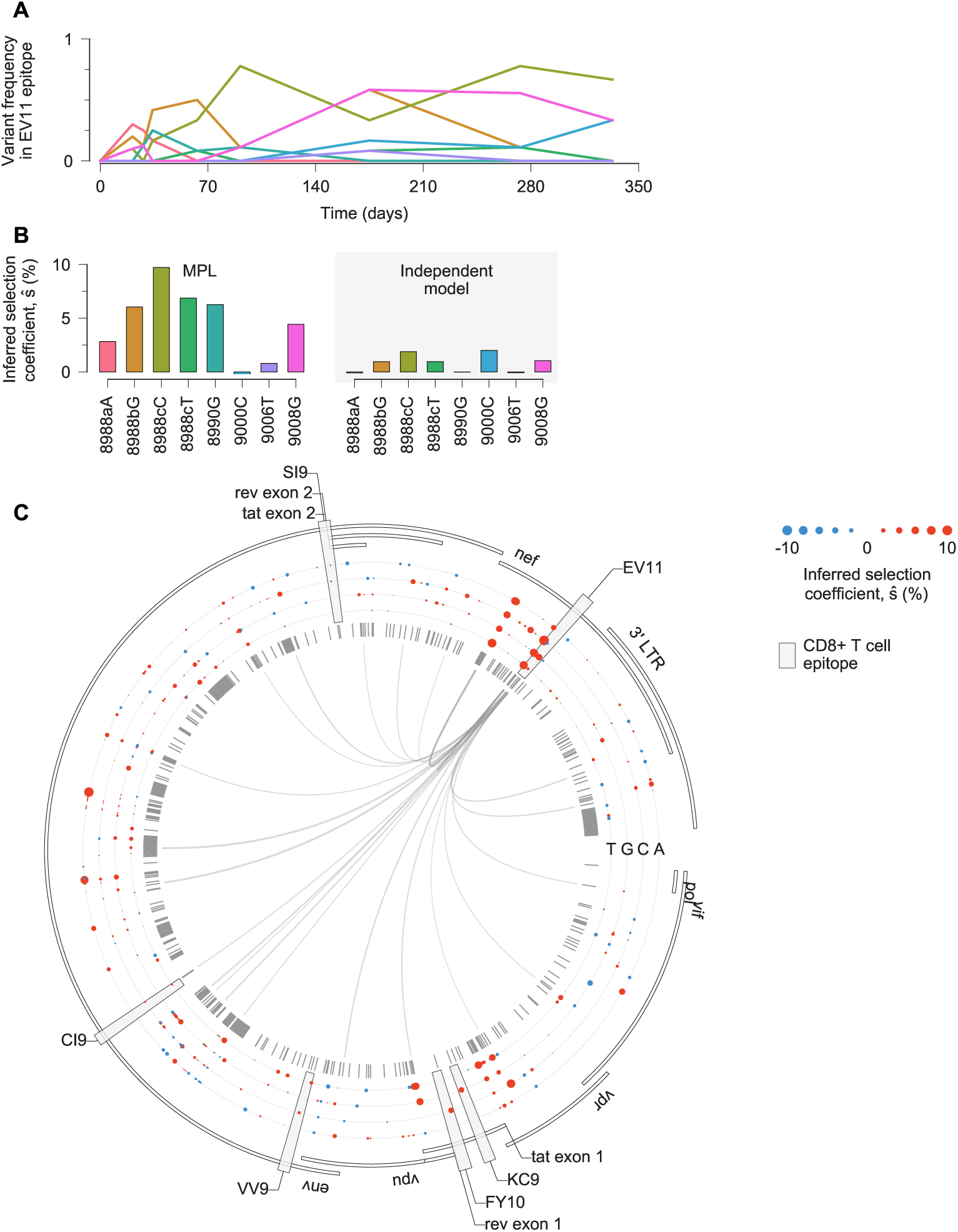
Estimates of selection coefficients in a complex example of clonal interference. **A**, Multiple escape variants for the Nef epitope EV11, targeted by individual CH131, interfere with one another over the course of nearly two years. Here we have omitted the trajectories for transient variants with a deletion at sites 8988a-8988c, which are insertions with respect to the HXB2 reference sequence. **B**, MPL infers that all nonsynonymous EV11 escape variants are positively selected. Variants 9000C and 9006T are both synonymous, and are inferred to be nearly neutral by MPL. As in previous examples, inferences using only the trajectories of individual variants only infer substantial positive selection for variants that are polymorphic at the final time point, or where the transmitted/founder (TF) allele at the same site appears strongly selected against. In the latter case, positive selection is inferred because all selection coefficients are normalized such that the selection coefficient for the TF variant is zero. This is why the independent model infers 8988T to be beneficial despite its low frequency at the final time point. Note that the independent model also infers one of the synonymous mutations to be beneficial. **C**, Inferred selection in the HIV-1 3′half-genome sequence for CH131. Inferred selection coefficients are plotted in tracks. Coefficients of TF nucleotides are normalized to zero. Tick marks denote polymorphic sites. Inner links, shown for sites connected to the EV11 epitope, have widths proportional to matrix elements of the inverse of the integrated covariance. Linked sites affect selection estimates within the epitope.

**Supplementary Fig. 9.**
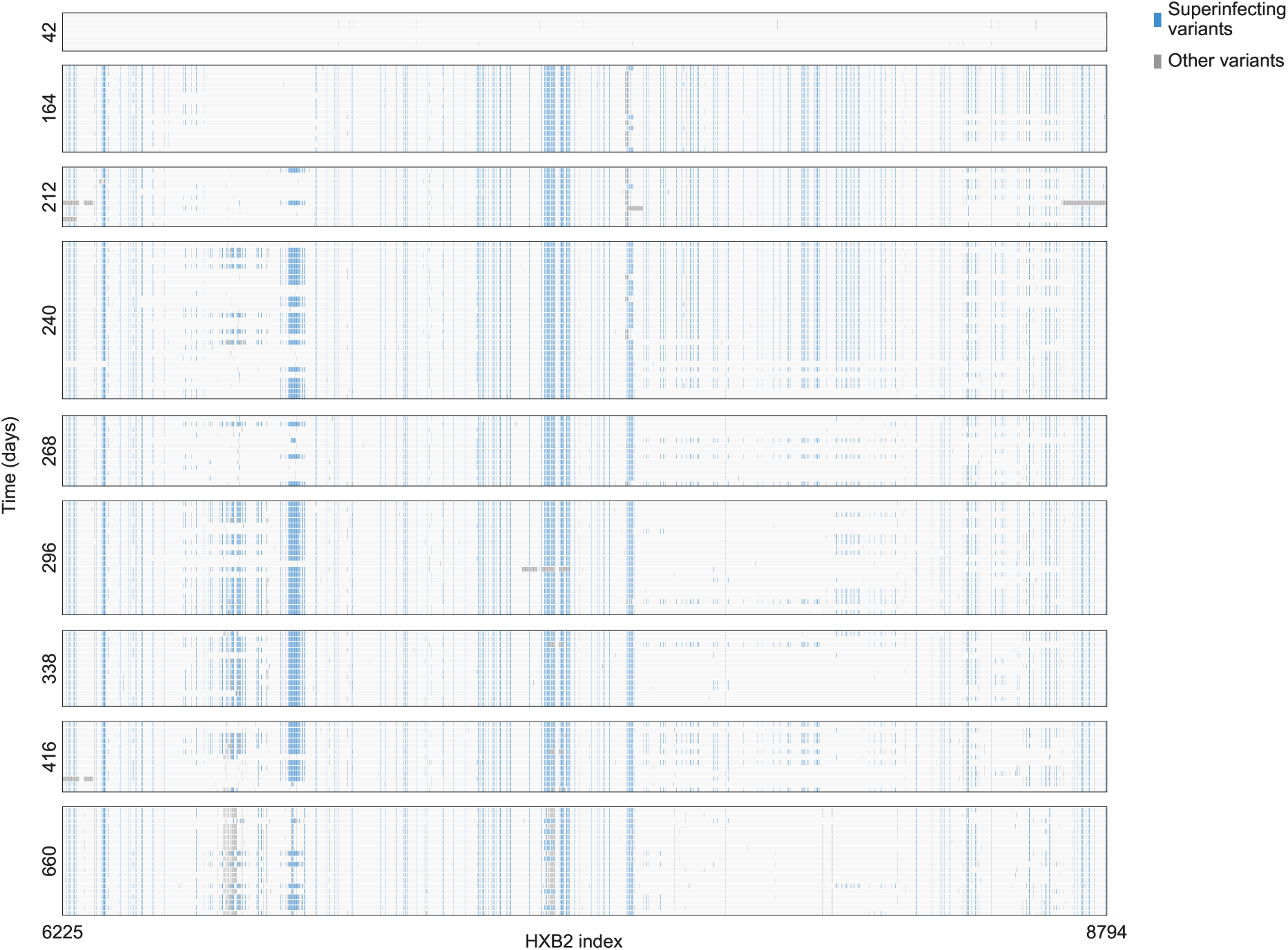
Extensive recombination between CAP256 primary and superinfecting strains. Individual CAP256 was infected by a distinct superinfecting strain of HIV-1 15 weeks after the primary infection (45). Each row represents a sequence, with nucleotide variants different from the primary infecting strain highlighted. Soon after superinfection, by 164 days after initial infection, recombinants between the primary and superinfecting strains dominate the viral population. Recombination continues throughout infection, introducing substantial variation into the VRC26 epitope region by 240 days after initial infection.

## Supplementary Information

### 1. Derivation of the Wright-Fisher path integral

This section presents the steps used to derive the path integral approximation for the probability of the mutant allele frequencies following a path (***x***(1), ***x***(2), *…*, ***x***(*T*)) conditioned on ***x***(0), as described in Methods. While we recall that the result was presented for a biallelic model with symmetric mutation probabilities, the technique is by no means limited by this model and can be extended to incorporate other model complexities, some of which are considered subsequently.

To summarize Section 1, we first describe the WF model in Section 1.1 in more detail, followed by presenting conditional moment expansions of the mean frequency vector in Section 1.2. These expansions will then be used in Section 1.3 to derive (S15), the path integral expression for the probability of a path of *genotype* frequencies (***z***(1), ***z***(2), *…*, ***z***(*T*)), conditioned on ***z***(0). In Section 1.4, this genotype-level path integral will be used to derive (S22), the path integral expression for the probability of a path of *allele* frequencies (***x***(1), ***x***(2), *…*, ***x***(*T*)), conditioned on ***x***(0).

#### 1.1. Wright Fisher model

We start with some brief comments on the WF model. At generation *t*, denote ***Z***(*t*) = (*Z*_1_(*t*), *…*, *Z*_*M*_ (*t*)) as the random genotype frequency vector, and thus ***z***(*t*) = (*z*_1_(*t*), *…*, *z*_*M*_ (*t*)) corresponds to an observed realization of this random vector. (These vectors, as well as all other vectors that we define, are taken as column vectors.) The stochastic dynamics of the genotype frequencies are governed by the transition probabilities reported in (2)-(4) of Methods. At generation *t* + 1, *y*_*a*_(*t*) recombination occurs, and as a consequence of this action, the mean frequency of genotype *a*, conditioned on ***Z***(*t*) = ***z***(*t*), is given by

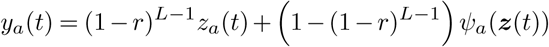

where the factor (1–*r*)^*L−*1^represents the probability of an individual not undergoing recombination, and *Ψ*_*a*_ (***z***(*t*)) the probability of forming genotype *a* after recombination from individuals in generation *t*. This latter quantity is given by

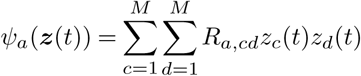

with *R*_*a,cd*_ denoting the probability that genotypes *c* and *d* recombine to form genotype *a*. The probability *R*_*a,cd*_ is a complicated function of the number of breakpoints and the particular genotypes *a, c* and *d*; however as we will show, we do not require the exact form of *R*_*a,cd*_ for our derivations. After recombination, the mean genotype frequencies at generation *t* + 1 are further shaped through selection and mutation. Specifically, after recombination, selection and mutation, the mean frequency of genotype *a*, conditioned on ***Z***(*t*) = ***z***(*t*), admits

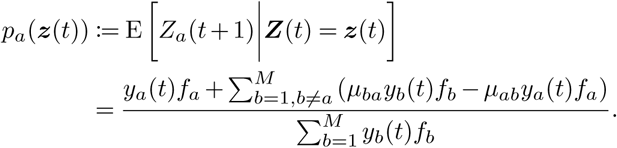

Then, the WF dynamics, specified by the transition probability

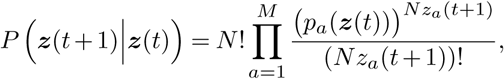

simply results from performing random multinomial sampling (of *N* individuals) with these mean frequencies.

Further, we recall the assumption that 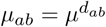, with *d*_*ab*_ the Hamming distance between genotypes *a* and *b*, and the additive model for genotype fitness, such that the selection coefficient of genotype *a* admits

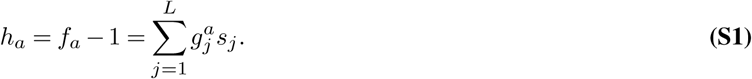

We may then write

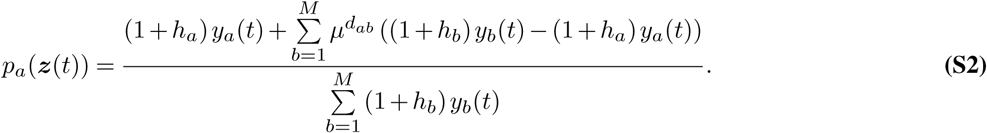

We work under the assumption that the population size *N* is large, and that as *N* → ∞,

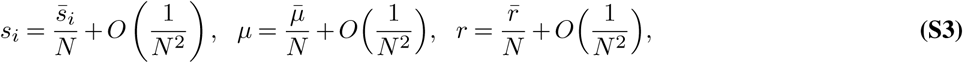

and consequently

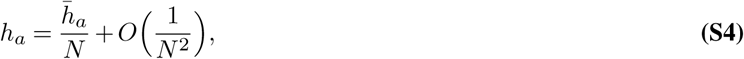

where 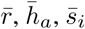 and 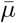 are constants that are independent of *N*.

#### 1.2. Conditional moment expansions

The diffusion approximation, and the path integral expression, depends on the asymptotic properties of the mean frequency vector *p*_*a*_(***z***(*t*)), as well as those of higher order moments. For large *N*, under the parameter scalings above,

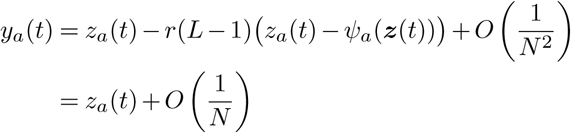

and thus (S2) can be expressed as

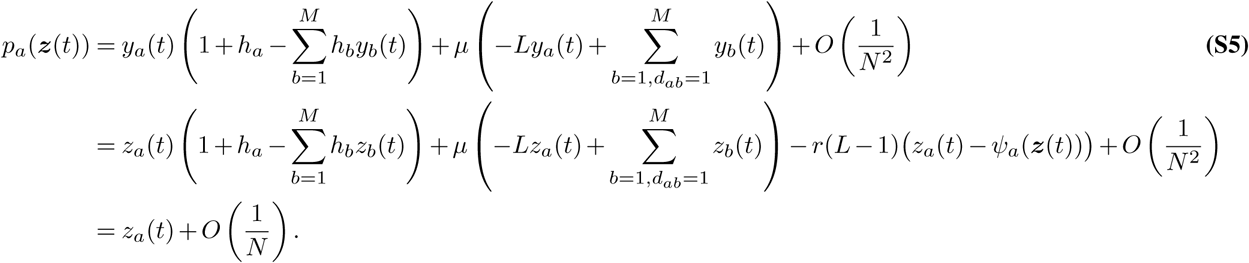

The conditional covariance of the genotype frequencies can also be expressed as

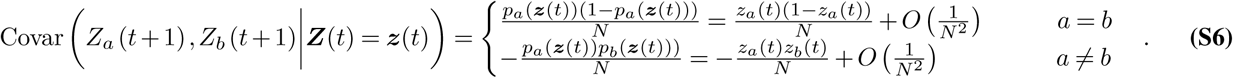

These expansions will be used in the following subsection.

For the subsequent analysis with incomplete temporal sampling (Section 2), we will also require the following expansions:

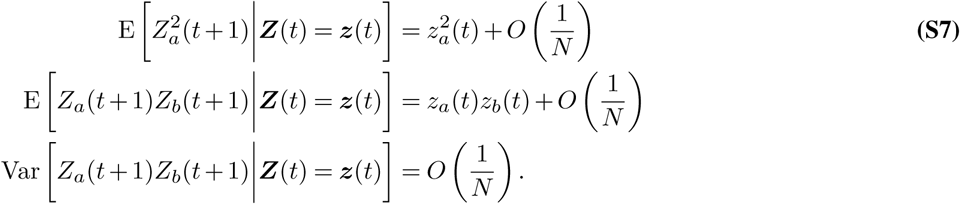

These results are easily obtained by computing the moment generating function of ***Z***(*t* + 1) conditioned on ***Z***(*t*) = ***z***(*t*),

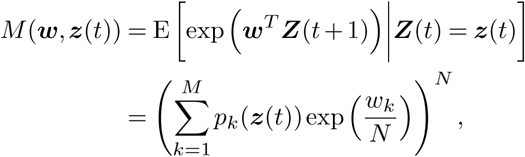

where ***w*** = (*w*_1_, *…*, *w*_*M*_), and then evaluating relevant derivatives at zero in the standard way.

#### 1.3. Genotype-level path integral

The moment expansions above give rise to a diffusion approximation and subsequently a path integral representation for the probability of genotype frequencies following a particular trajectory. The diffusion approximation is a continuous-time continuous-frequency approximation to the discrete-time discrete-frequency WF process. It is valid under the large-*N* parameter scalings (S3) and (S4), and corresponds to the continuous process

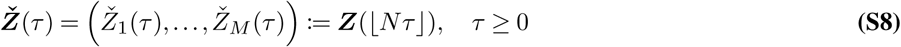

taken in the limit *N*→ ∞, where ⌊.⌋ denotes the floor function. Here *τ* is a continuous time variable with units of *N* generations, with one generation in discrete time (i.e., from *t* to *t* + 1) thus taking

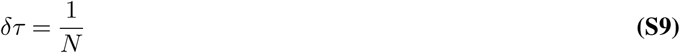

continuous time units. The diffusion process is described by the probability density function *ϕ*, the solution to

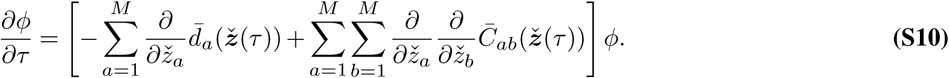

This is characterized by the drift vector 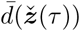, which describes the rate of expected changes in genotype frequencies at time *τ*, and the diffusion matrix 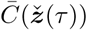, which describes the scaled covariance of the genotype frequency changes.

The drift vector has *a*th entry (see equation 4.99 of Risken(1))

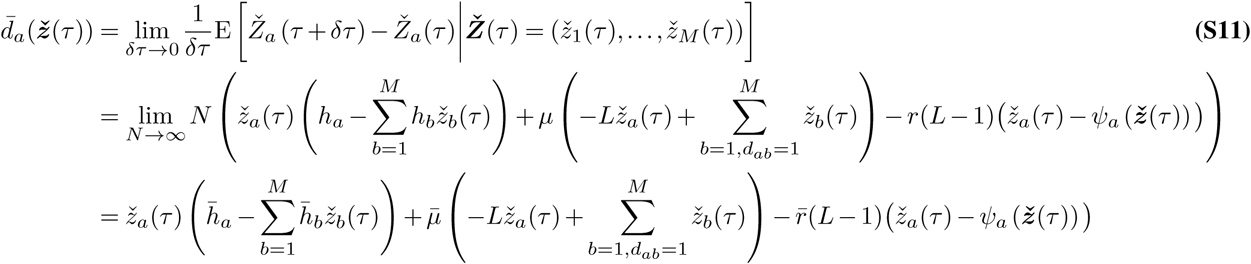

where the second line follows from (S9), (S8), and (S5). (As a technical point, we note that upon making the replacement *t* = ⌊ *Nτ* ⌋, the variable *t* is considered fixed when taking limits over *N*. That is, we may write

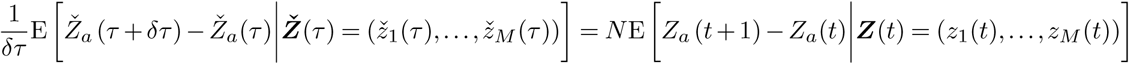

and the limit of the left-hand side as *δτ* →0 coincides with that of the right-hand side as *N*→ ∞. The same arguments will also be employed, although not explicitly stated, when taking limits in the subsequent derivations of drift vectors and diffusion matrices.) The diffusion matrix has (*a, b*)th entry (see equation 4.100 of Risken(1))

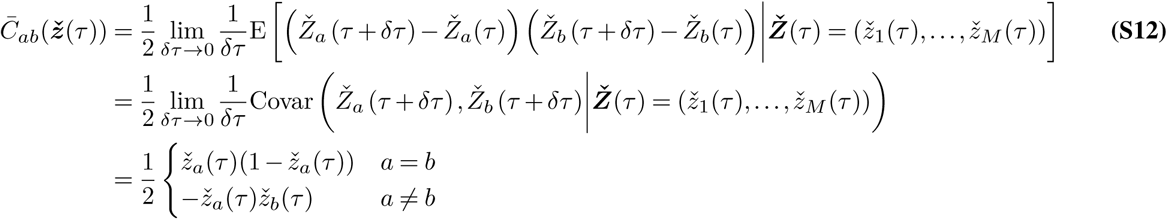

where the second line follows from noting that

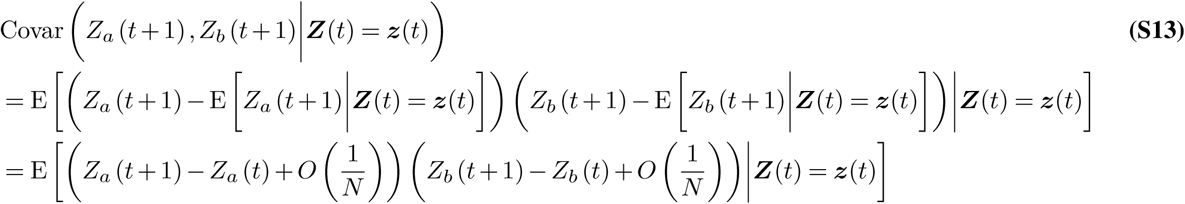

along with (S8) and (S9), while the last line of (S12) follows from (S6).

The path integral (see equation 4.109 of Risken(1)) approximates the transition density of a diffusion process over a small time period. Specifically, applying this approach to the process described by (S10), for small *δτ* (equivalently large *N*), we obtain for the transition probability density over a single generation,

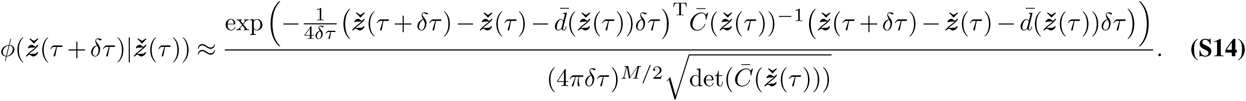

From this result, the transition probability for a single generation of the original discrete-time discrete-frequency WF process can (for large *N*) be approximated by

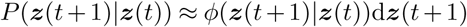

where the 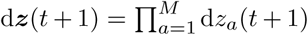 represent small frequency differences accounting for the quantization of the continuous genotype frequency space at each time point. The probability of observing a trajectory of genotype frequencies (***z***(1), ***z***(2),*…*, ***z***(*T*)) is then given by

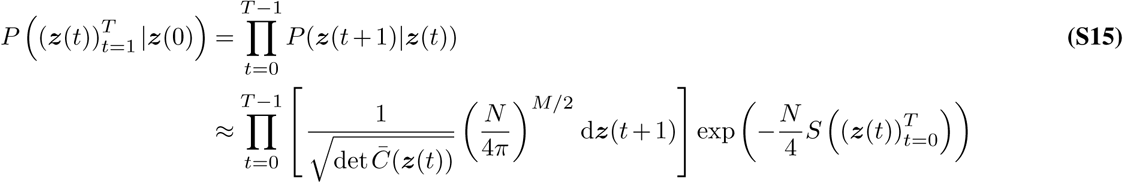

where

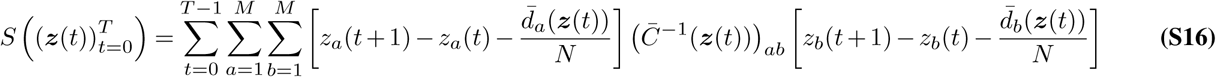

which is the desired path integral expression. In the language of physics, 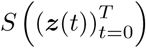 is referred to as the *action*.

#### 1.4. Mutant allele-level path integral

Based on the genotype transition probability density (S14), we now provide an approximation for the mutant allele transition probability density. Let ***x***(*t*) = (*x*_1_(*t*),*…, x*_*L*_(*t*)), for which

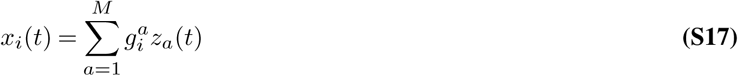

is the mutant frequency at locus *i* during generation *t*. Also, define the random mutant allele frequency vector ***X*** (*t*) = (*X*_1_(*t*),*…, X*_*L*_(*t*)), which from (S17) is related to the random genotype frequency vector by

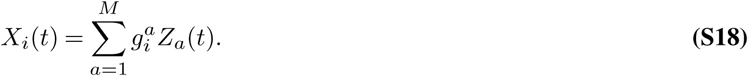

The observed frequency vector ***x***(*t*) is thus a realization of this random vector. The continuous process which characterizes the mutant allele frequencies is

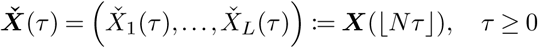

taken as *N* → ∞, which satisfies

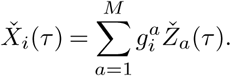

This allele frequency process is a diffusion process, whose probability density evolution is described by a solution to

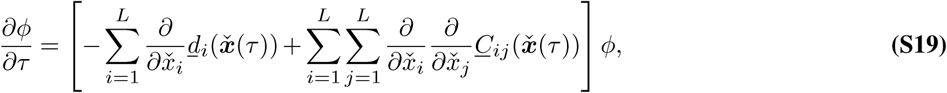

characterized by the drift vector 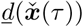 and diffusion matrix 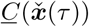. Note that the drift and the diffusion equations given in the Methods are the same as here in the Supplementary Information, but those in the Methods use simpler notation by not explicitly distinguishing between continuous-time and discrete-time processes.

In the genotype case, the transition probability density was approximated to have a Gaussian form (S14). As the mutant allele frequencies are linear combinations of the genotype frequencies (S17), this implies that the transition probability density of mutant alleles also has a Gaussian form. Therefore, the allele-level drift and diffusion terms in (S19) are also a linear combination of the genotype drift and diffusion terms. The drift vector thus has *i*th entry

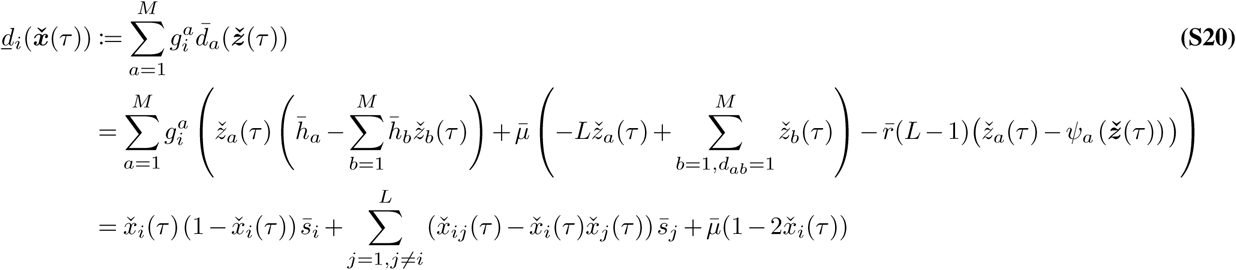

with

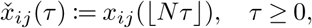

and the last line follows by applying (S1)

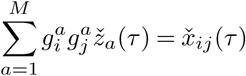

and by noting that 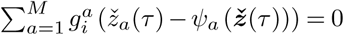. To see why the latter holds, first define

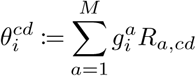

which is the probability that genotypes *c* and *d* recombine to form a genotype which has a mutation at locus *i*. Since the model is biallelic, the recombination event could involve allele pairs (0, 0), (0, 1), (1, 0), or (1, 1) at locus *i* of genotypes *c* and *d*. We thus have

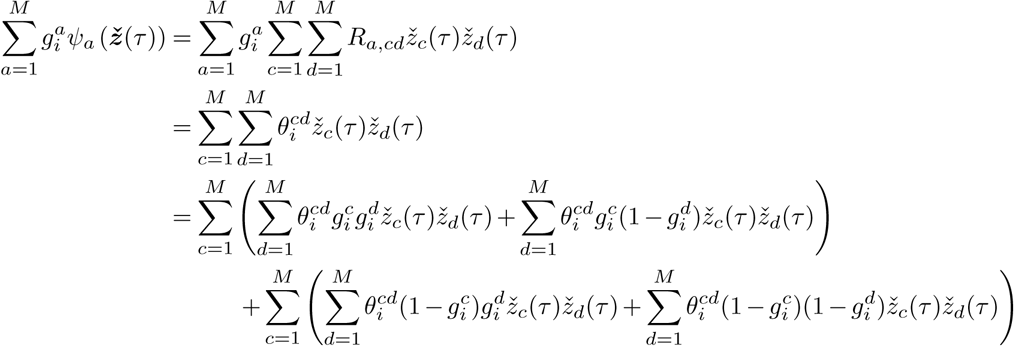

Now by noting that

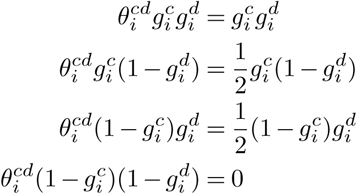

where the factor of 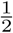 arises because there is a 50% chance that genotype *c* (*d*) with a mutant at locus *i* and genotype *d* (*c*) with WT at locus *i* will recombine to a genotype with a mutant at locus *i*, we have

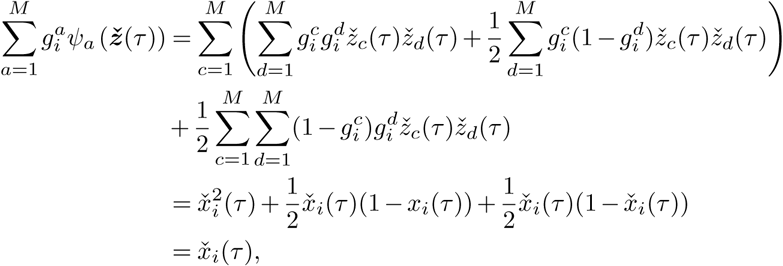

thus implying that.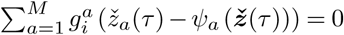.

(Note that above and also in the subsequent derivations, since we are focused on the mutant allele frequency dynamics, we adopt notation which only explicitly demonstrates dependencies on the mutant allele frequencies. It should be recognized, however, that these dynamics also depend on the pairwise mutational frequencies. We do not explicitly show this, for the sake of notational convenience.) Considering now the diffusion matrix, this has (*i, j*)th entry

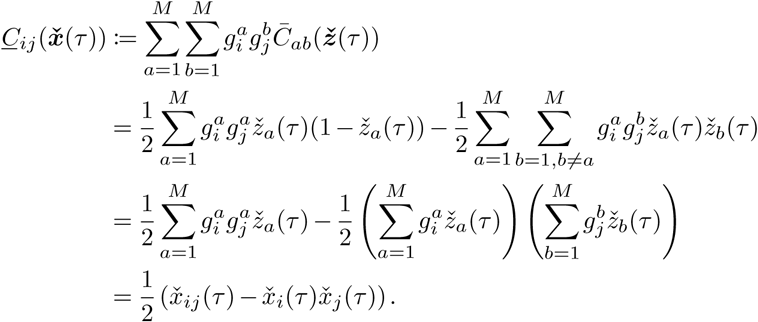

Note that if we concatenate the infinitesimal drift for all loci in vector form, it yields

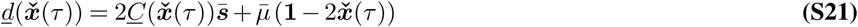

where **1** is the vector of ones.

As for the genotype case, equation 4.109 of Risken (1) may be applied, which approximates the transition probability density of this diffusion process over a single generation by a Gaussian distribution, given as

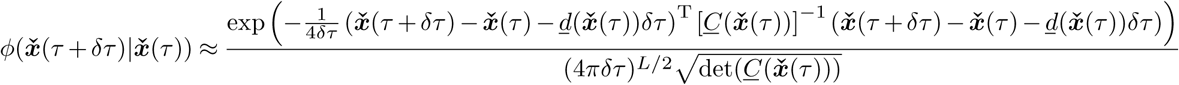

From this result, and recalling that *δτ* = 1*/N*, the transition probability for a single generation of the original discrete-time discrete-frequency WF process can (for large *N*) be approximated by

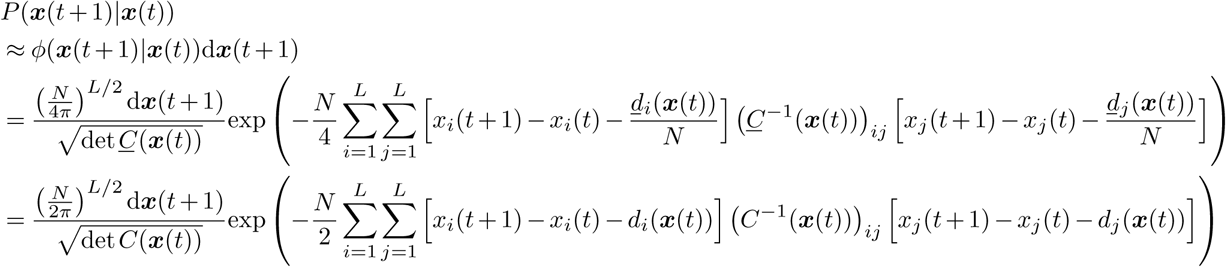

where the 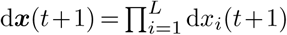 represents small frequency differences accounting for the quantization of the continuous mutant allele frequency space, and we have made the replacements *<u>d</u>*_*i*_(***x***(*t*)) = *Nd*_*i*_(***x***(*t*)) and (*<u>C</u>*(***x***(*t*)))_*ij*_ = (1*/*2)*C*(***x***(*t*))_*ij*_. The path integral expression then follows by noting that the probability of observing a trajectory of mutant allele frequencies (*x*(1),*x*(2), …, *x*(*T*)) is given by

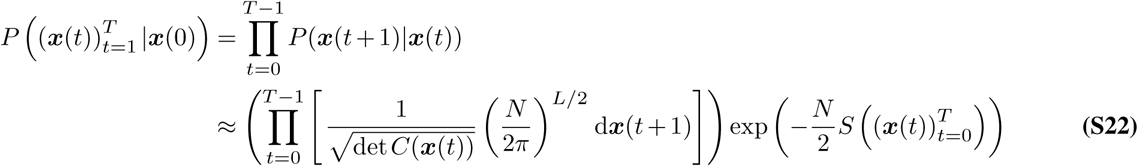

where

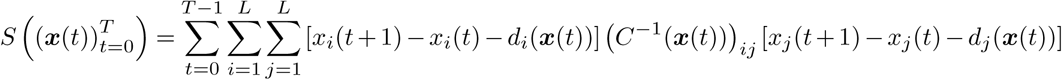

As in (S16),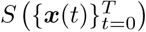 is referred to as the action in physics.

### 2. Derivation of the MPL estimator of allele selection coefficients

This Section first presents a proof of the MPL estimate (10) of the selection coefficients. As stipulated in Methods, this is obtained as the MAP estimate of the selection coefficients, which is the solution to

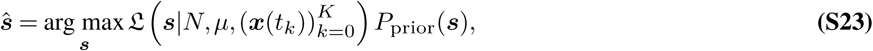

where

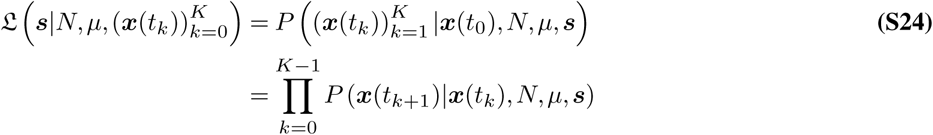

is the likelihood function, while (***x***(*t*_0_), ***x***(*t*_1_), *…*, ***x***(*t*_*K*_)) is the observed trajectory of mutant allele frequencies.

The main challenge in solving (S23) is that it requires computing the likelihood (S24), which is complicated. This is simplified by adopting the path integral approach outlined in Section 1; but now extending the analysis to account for sampling times *t*_0_, *…*, *t*_*K*_ spanning multiple generations. Other than this difference, we may follow the same approach, starting by giving asymptotic moment expansions in Section 2.1. These expansions will then be used in Section 2.2 to derive (S32), the probability of a path of *genotype* frequencies (***z***(*t*_1_), ***z***(*t*_2_), *…*, ***z***(*t*_*K*_)), conditioned on ***z***(*t*_0_), and subsequently (S33), the probability of a path of *allele* frequencies (***x***(*t*_1_), ***x***(*t*_2_), *…*, ***x***(*t*_*K*_)), conditioned on ***x***(*t*_0_). This allele-level path integral will then be used to derive the MPL estimate (10) in Section 2.3. We then show in Section 2.4 that this MPL estimate can also be derived based on the genotype-level path integral, demonstrating no loss of optimality from derivations based on an allele-level representation. Finally, we extend the biallelic and symmetric mutation modelling assumptions to account for multiple alleles per locus and asymmetric mutation probabilities in Section 2.5. This will be required to analyze the HIV data in the main text.

#### 2.1. Conditional moment expansions

The mean frequency of genotype *a* at generation *t* + Δ*t*, conditioned on ***Z***(*t*) = ***z***(*t*), admits

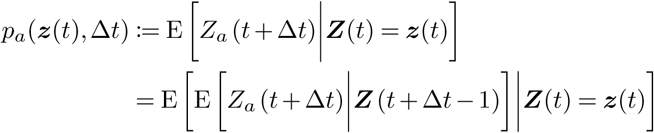

where the second line follows from the law of total expectation. Note that *p*_*a*_(***z***(*t*), 1) ≡ *p*_*a*_(***z***(*t*)), with *p*_*a*_(***z***(*t*)) defined in (S2). Now applying (S5), we further obtain

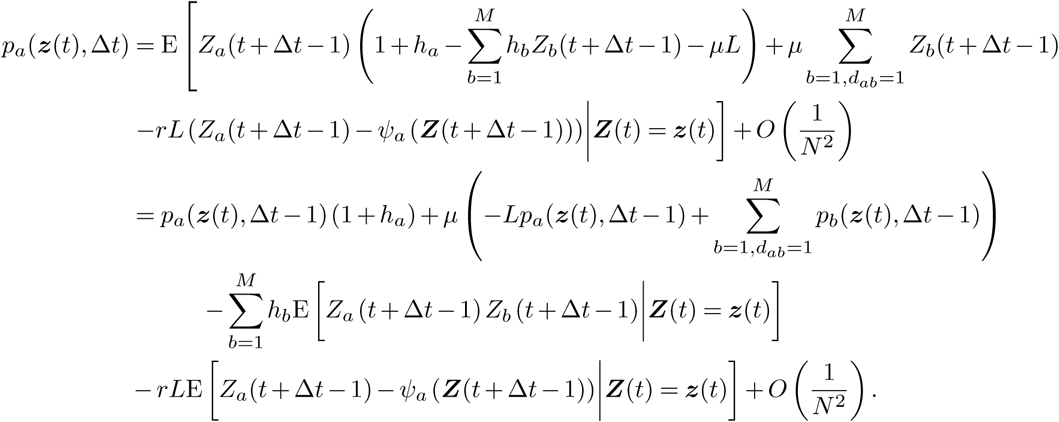

By iterating the above procedure Δ*t* times, applying the expansions in (S5) and (S7), and recalling (S3) and (S4), we arrive at the explicit expansion

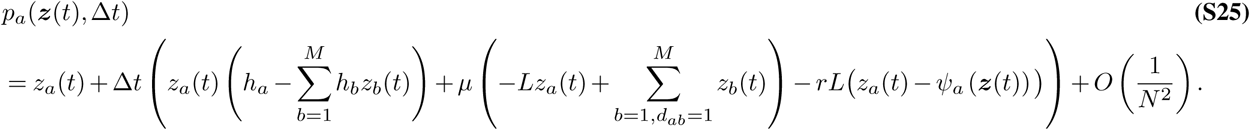

Next, we turn to deriving corresponding expansions for the conditional variance and covariance. Using the law of total variance, the conditional variance of the frequency of genotype *a* at generation *t* + Δ*t*, conditioned on ***Z***(*t*) = ***z***(*t*), is given by

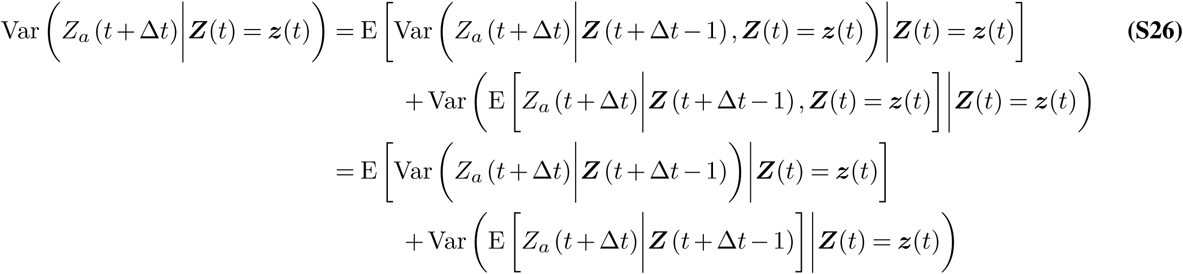

where the second line follows from the Markov property of the WF process. The first term in (S26) can be written as

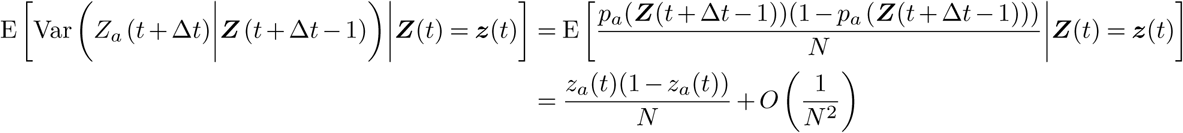

which was obtained by iteratively applying the expansions (S5) and (S7). Moreover, the second term in (S26) admits

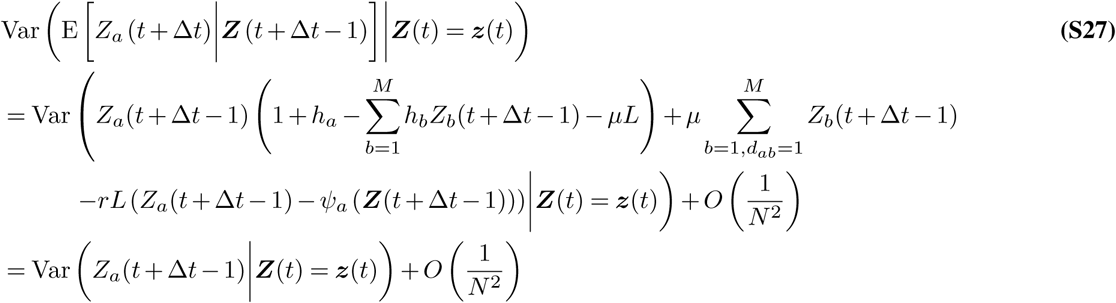

where the second line follows from (S5) and the last line follows from first noting that for arbitrary random variables 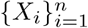, we have

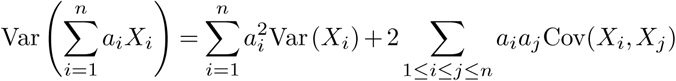

recalling again (S3) and (S4), and applying the expansions (S6) and (S7). Substituting (2.1) and (S27) into (S26), and iterating, we arrive at

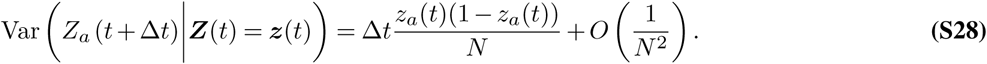

The expansion for the conditional covariance is obtained by following a similar procedure. For *a* ≠ *b*, it leads to

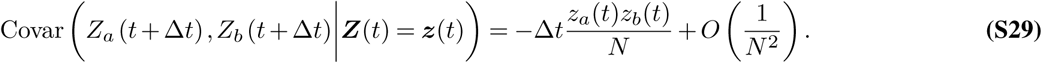

#### 2.2. Path integral representation of the likelihood function

Based on the above conditional moment expansions, for the observed set of time points *t*_0_ < *t*_1_ < …< *t*_*K*_, we can present a path integral expression for the likelihood function (S24). The proof follows a similar procedure to that in Sections 1.3 and 1.4, with basic modifications to the drift vector and covariance matrix to account for frequency vector observations being taken multiple generations apart.

First, we consider genotype frequency evolution, and assume that genotype frequencies are observed at generations *t* and *t* + Δ*t*, with no observations in-between. Then, after mapping from discrete-time to continuous-time and taking *N* → ∞ as before (i.e., in equation S8), the modified drift vector now characterizes the expected infinitesimal change in genotype frequencies between continuous time points *τ* and *τ* + Δ*tδτ*, and the covariance matrix characterizes the second moment of the change in genotype frequencies between the two time points. Specifically, using (S25), the *a*th element of the modified drift vector can be written as (see equation 4.99 of Risken(1))

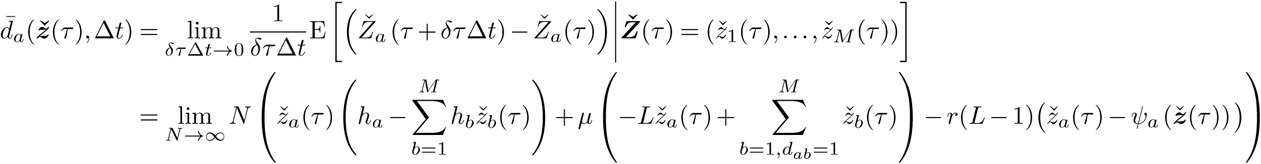

which turns out to be equivalent to (S11), and hence

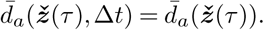

An analogous equivalence also holds for the modified diffusion matrix. Specifically, this has (*a, b*)th entry (see equation 4.100 of Risken (1))

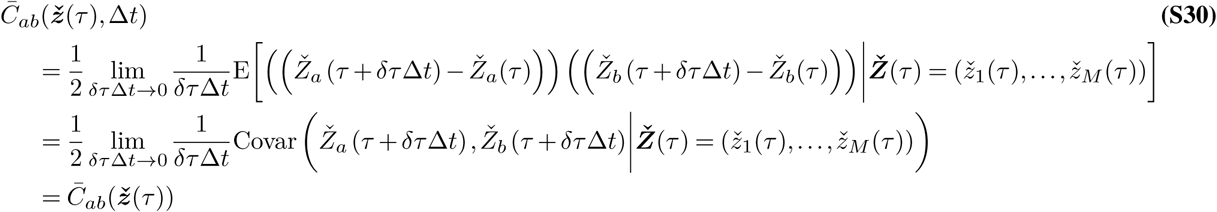

where the second line follows via the same arguments as in (S13), while the last line was obtained by applying the expansions (S28) and (S29).

As before, the transition density can then be approximated for small *δτ* Δ*t* as

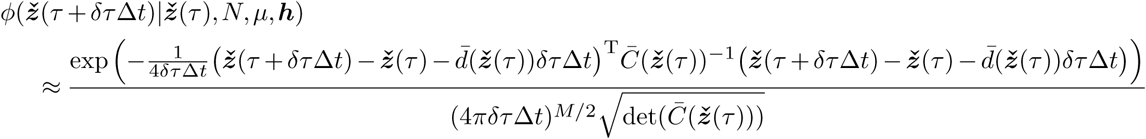

Here we show explicit dependence on *N*, *μ* and ***h***. From this result, and recalling that *δτ* = 1*/N*, the transition probability for a single generation of the original discrete-time discrete-frequency WF process can (for large *N*) be approximated by

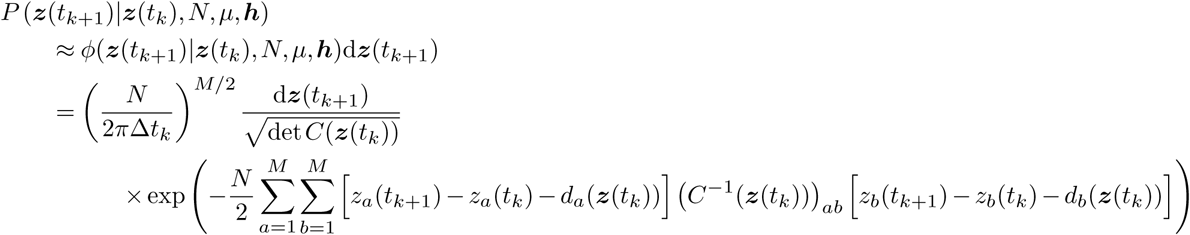

where Δ*t*_*k*_ = *t*_*k*+1_ − *t*_*k*_, the 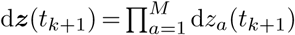 represent small frequency differences accounting for the quantization of the continuous genotype frequency space, and where we have defined 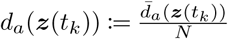 and

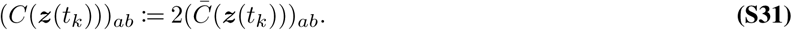

The path integral expression then follows by noting that the probability of observing a trajectory of genotype frequencies (***z***(*t*_1_), ***z***(*t*_2_), …, ***z***(*t*_*K*_)) is given by

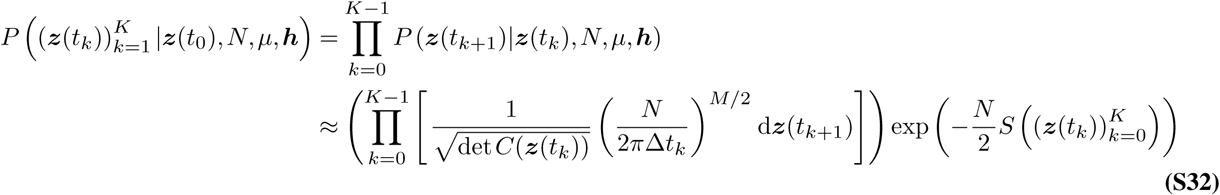

where

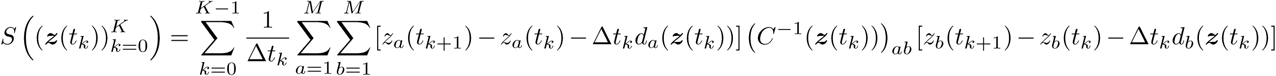

Based on these results, a corresponding path integral approximation for the probability of observing a trajectory of mutant allele frequencies (***x***(*t*_1_), ***x***(*t*_2_), …, ***x***(*t*_*K*_)), and hence for the likelihood function (S24), is then obtained by mirroring the steps in Section 1.4, giving

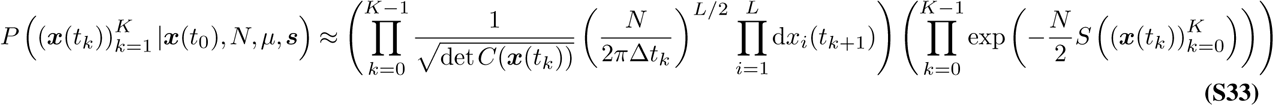

where

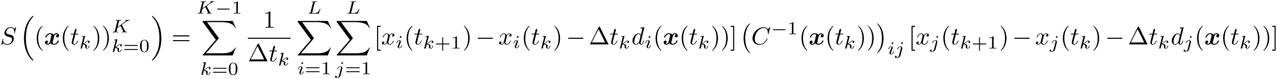

#### 2.3. The MPL estimator solution

Returning to the MAP problem (S23), we note that this is equivalent to

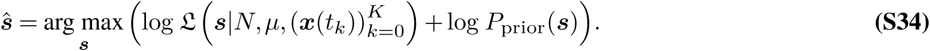

The likelihood 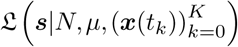 is given by (S24) and is approximated (S33). Assuming a conjugate-prior distribution

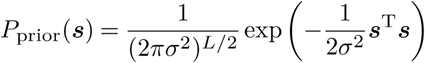

we take the vector derivative of the right-hand side of (S34) with respect to ***s*** (see equation 10, Chapter 10.2.1 of Lutkepohl(2)), equate to zero, and then solve for ***s***. This yields

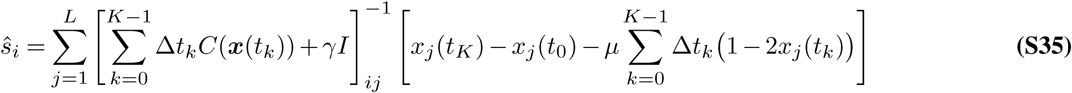

for *i* = 1, …, *L*, where *γ* = 1*/Nσ*^2^. This is the desired *MPL estimate* of the selection coefficients, reported in (10) of Methods. Here, we note that *γ* can also be viewed as a parameter that regularizes the covariance matrix prior to inversion.

#### 2.4. Equivalence of genotype- and allele-level analyses

While the MPL estimator was derived using a path integral expression for the probability of mutant allele frequency trajectories, the WF evolutionary process is defined for genotypes. Hence, one may naturally ask whether there is any loss in information in considering only the marginal frequency dynamics. As we now show, the answer is no. Specifically, we show that the estimate of the selection coefficients based on genotype frequencies is equivalent to the derived MPL estimate.

We begin by defining *G* as a *M* × *L* matrix with (*a, j*)th entry 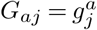 Let ***h*** = (*h*_1_, *h*_2_, …, *h*_*M*_) denote the column vector of genotype selection coefficients, which relates to the mutant allele selection coefficients via

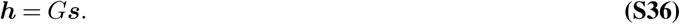

Given the observed genotype frequencies (***z***(*t*_0_), ***z***(*t*_1_), …, ***z***(*t*_*K*_)), the MAP estimator of the selection coefficients admits

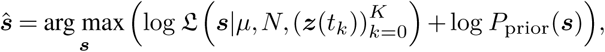

where the likelihood function is given as

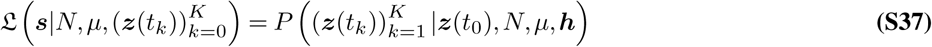

Note that the right-hand side of (S37) is approximated by (S32). Expressing the genotype selection coefficients in (S32) in terms of mutant allele selection coefficients using (S36) gives the likelihood of the mutant allele selection coefficients ***s*** for an observed genotype frequency path and parameters *N*, *μ*. Differentiating the resulting expression with respect to ***s*** and equating to zero leads to

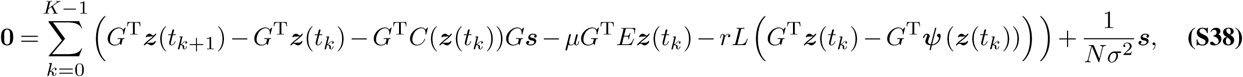

where 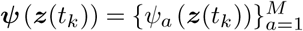 and *C*(***z***(*t*_*k*_)) is the genotype covariance matrix as given by (S30) and (S31), i.e., with elements

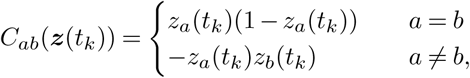

while matrix *E* has elements

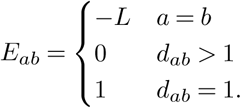

Noting that the relation between allele and genotype frequencies (S18) can be expressed in vector form as ***x***(*t*_*k*_) = *G*^T^***z***(*t*_*k*_), and that *C*(***x***(*t*_*k*_)) = *G*^T^*C*(***z***(*t*_*k*_))*G* and *G*^T^***Ψ*** (***z***(*t*_*k*_)) = ***x***(*t*_*k*_), we can solve (S38) to obtain the MAP estimate of the allele selection coefficients

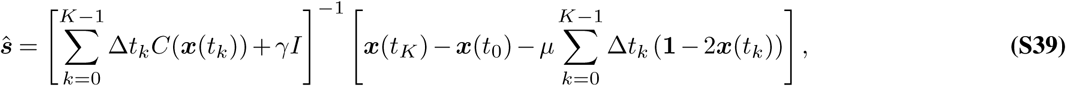

where *γ* = 1*/Nσ*^2^. This is the same as the MPL estimator (S35).

This equivalence is important. It implies that under the additive fitness model, only the single and pairwise mutational frequencies are needed for optimally estimating the selection coefficients, and higher order information (e.g., three-locus mutational frequencies) are irrelevant. The same equivalences can also be shown for extensions of the MPL estimator presented in the following section. Further extensions will be explored in future work.

#### 2.5. Extension to multiple alleles per locus and asymmetric mutation probabilities

Here we demonstrate the extension of our inference framework to incorporate multiple alleles per locus and asymmetric mutation probabilities. The same notation and definitions introduced in Section 1 and 2 will also be used, unless stated otherwise, but will be interpreted in terms of the extended model. For example, ***Ž*** (*τ*) was introduced in (S8) to refer to the binary genotype continuous process with symmetric mutation probabilities. This notation will also be used here, but it will now refer to the non-binary genotype continuous process with asymmetric mutation probabilities. We first describe the model in detail.

We assume there are ℓ alleles per locus, thus resulting in *M* = ℓ^*L*^ genotypes, with the multi-allelic sequence for genotype *a* denoted by 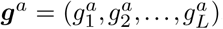, where 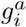 is the allele at locus *i*. Nucleotide sequences, for example, will have 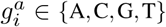. For convenience, we denote each allele with an integer representation, and consider the fitness of each genotype with respect to a reference sequence comprising of allele ℓ at each locus, i.e., the reference sequence is a sequence of ℓs. For example, if ℓ = 4 and *L* = 3, then the reference sequence is represented by (4, 4, 4). We assume, once again, an additive model of fitness, and that the reference genotype has a fitness of one. Under these assumptions, the selection coefficient for genotype *a* is given by

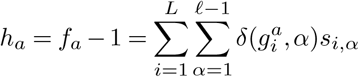

where *s*_*i, α*_ is the selection coefficient of allele *α* at locus *i*, and where *δ*(·, ·) is the Kronecker-delta function,

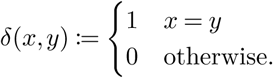

Note that the assumptions above imply that *s*_*i*,ℓ_ = 0 for all *i* = 1, …, *L*.

The allele frequency vector, describing the frequency of all alleles except (the reference) allele one, is given by ***x***(*t*) = (*x*_1,1_(*t*), …, *x*_1,ℓ−1_(*t*), …, *x*_*L*,1_(*t*), …, *x*_*L*,ℓ−1_(*t*)), where *x*_*i,α*_(*t*) denotes the observed frequency of allele *α* at locus *i* during generation *t*, and is related to the genotype frequencies by

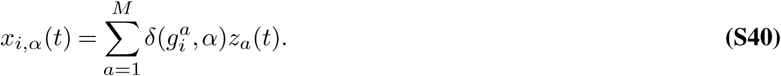

Finally, we denote *μ*_*αβ*_ as the mutation probability per generation from allele _*α*_ to allele *β* and *μ*_*ab*_ as the mutation probability per generation from genotype *a* to genotype *b*. These are related by

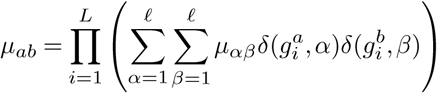

Given this model, the MAP estimate of the selection coefficients is the solution to

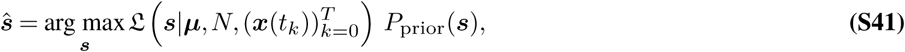

where

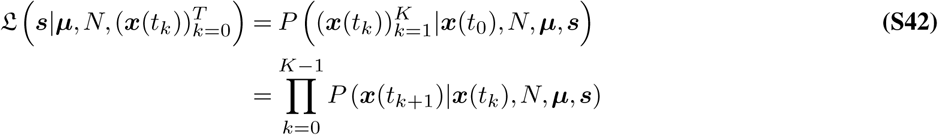

is the likelihood of the selection coefficients ***s*** = (*s*_1,1_, …, *s*_1,ℓ−1_, …, *s*_*L*,1_, …, *s*_*L*,ℓ−1_) and *P*_prior_(***s***) their prior distribution.

As with the simpler biallelic and symmetric mutation probability scenario, the main challenge in solving (S41) is that it requires computing the likelihood (S42), which is complicated. This is simplified as before, by adopting the path integral approach outlined in Section 1; but now extending the analysis to account for sampling times spanning multiple generations, multiple alleles per locus and asymmetric mutation probabilities. Other than these differences, we may follow the same approach, starting by giving asymptotic moment expansions. By direct analogy to (S3) and (S4), we will work under the assumption that *N* is large, and that as *N* → ∞,

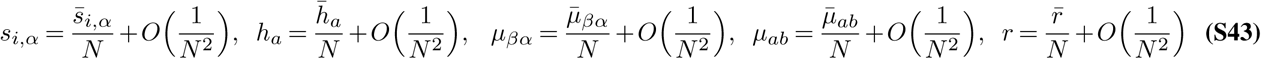

where 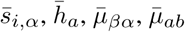 and 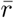 are constants independent of *N*.

##### 2.5.1. Conditional moment expansions

The mean frequency of genotype *a* at generation *t* + 1, conditioned on ***Z***(*t*) = ***z***(*t*), admits

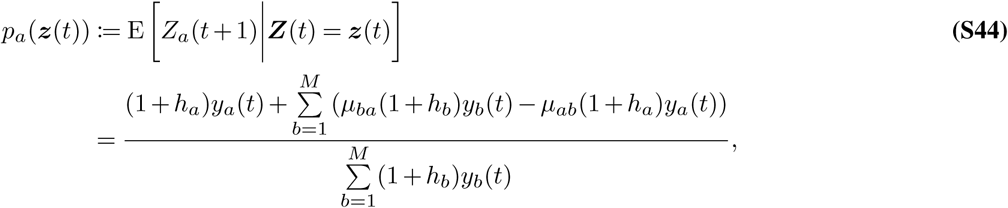

where recall that

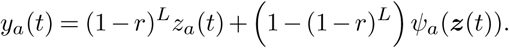

Utilizing (S44) along with (S43), and applying a similar proof to that described in Section 2.1, the mean frequency of genotype *a* at generation *t* + Δ*t*, conditioned on ***Z***(*t*) = ***z***(*t*), then admits

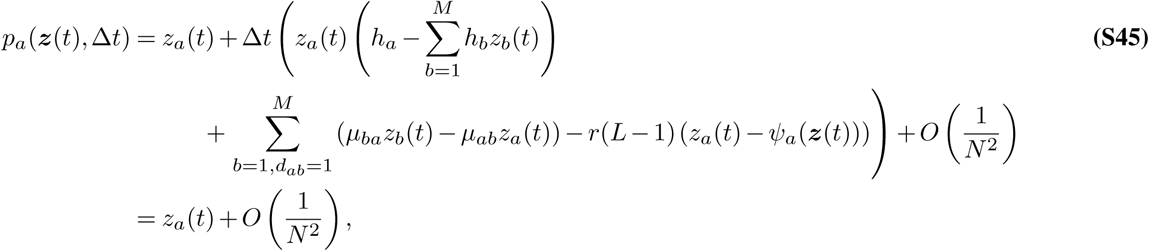

where *d*_*ab*_ here denotes the number of loci for which the allele identity at genotypes *a* and *b* differ, i.e.,

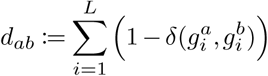

The corresponding expressions for the variance and covariance of the frequency of genotype *a* at generation *t* + Δ*t*, conditioned on ***Z***(*t*) = ***z***(*t*) are obtained analogously to (S28) and (S29) as

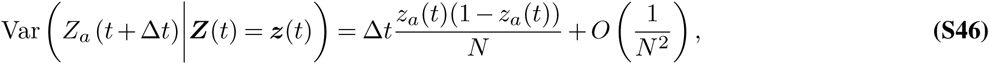

and

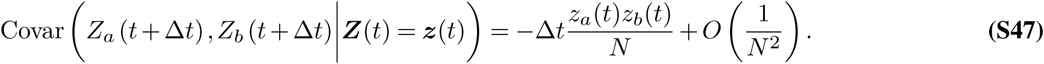

##### 2.5.2. Path integral representation of the likelihood function

Based on the above conditional moment expansions, we can derive a path integral representation for the likelihood function (S42). Starting by considering genotype evolution, the drift vector and diffusion matrix in this case are given respectively by

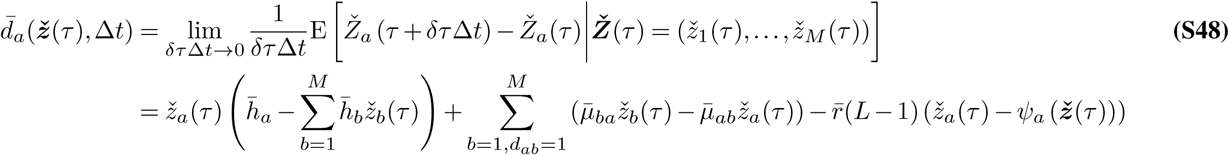

and

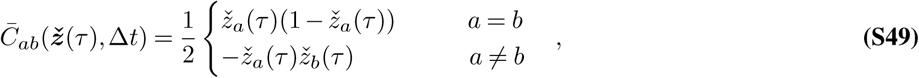

which follow from (S45), (S46), (S47), and by adopting the procedure described in Section 2.2. From these results, and again following the procedure in Section 2.2, the conditional probability of observing a trajectory of genotype frequencies (***z***(*t*_1_), ***z***(*t*_2_), …, ***z***(*t*_*K*_)) is obtained as

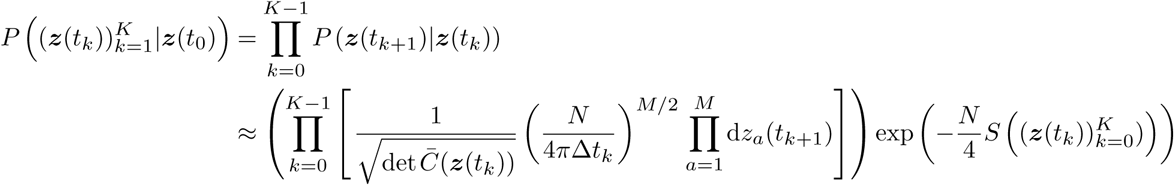

where Δ*t*_*k*_ = *t*_*k*+1_ − *t*_*k*_ and

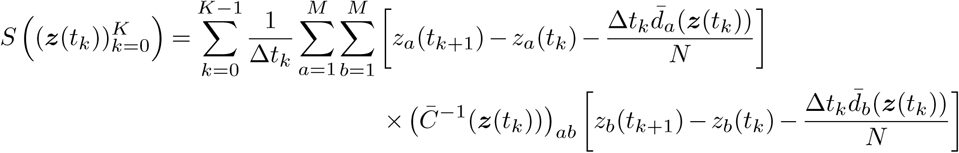

In this last equation we have dropped the second argument of the drift vector and diffusion matrix, as both quantities turn out to be independent of Δ*t*, as seen from (S48) and (S49).

Now turn to mutant allele evolution. By noting the linearity of expectation and the genotype-to-allele mapping (S40), the mean frequency of allele *α* at locus *i* during time *τ* + *δτ* Δ*t*, conditioned on 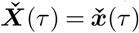, is given by

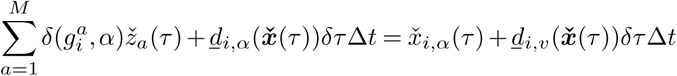

where

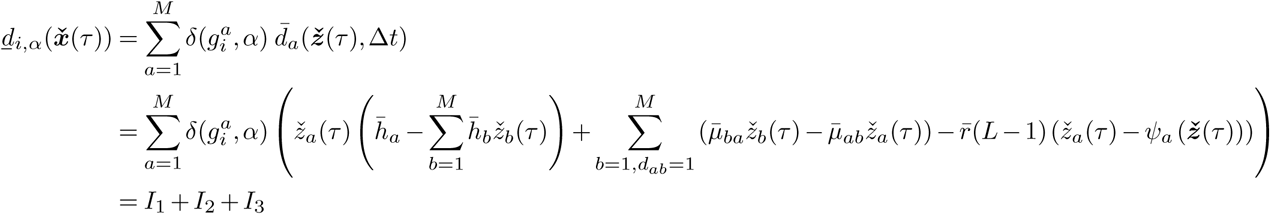

with the second line following from (S48). For *I*_1_, we have

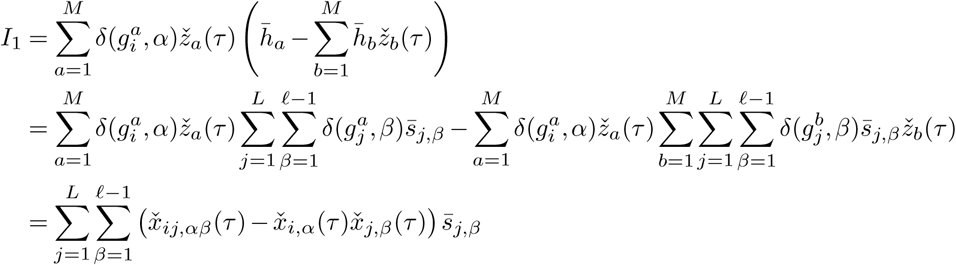

where 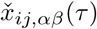 denotes the frequency of allele *α* and *β* occurring respectively at loci *i* and *j* during time *τ*, and given by

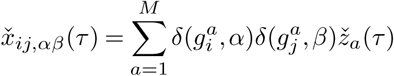

For *I*_2_, we have

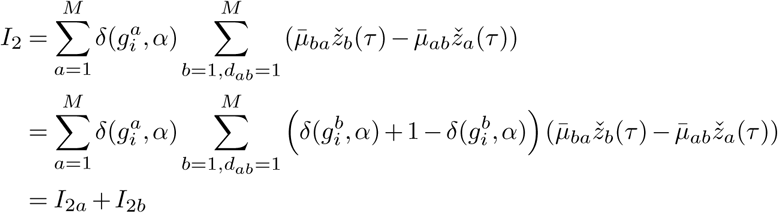

where

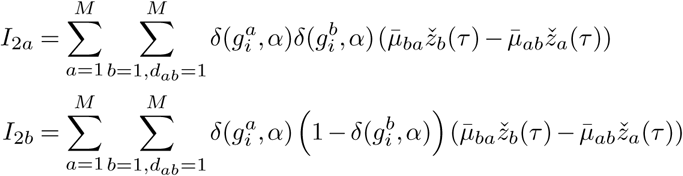

Consider now *I*_2*a*_, where we observe that 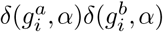 is non-zero only when genotypes *a* and *b* both have allele *α* at locus *i*, while (ii) the summation 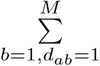 is over all genotypes *b* which have a different allele from genotype *a* at only one locus. These two observations imply that 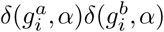 is non-zero, for *b* = 1, …, *M*, with *d*_*ab*_ = 1, only if *a* and *b* have a different allele at a single locus, but where the position of this locus is different from *i*. To illustrate this, consider a simple example with *L* = 2 and ℓ = 3, in which case

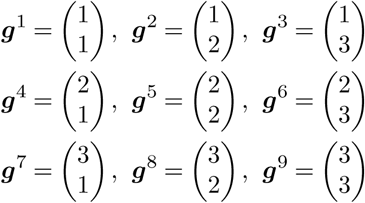

Then if *i* = 1 and *α* = 1, the only genotype-pairs (*a, b*) which result in a non-zero 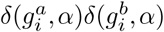 (subject to the condition *d*_*ab*_ = 1) are (1, 2), (2, 1), (1, 3), (3, 1), (2, 3) and (3, 2), as these genotype-pairs differ only at the second locus, while having allele one at locus *i* = 1.

Observe that every genotype-pair which result in a non-zero 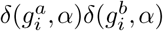 (subject to the condition *d*_*ab*_ = 1) occur in conjugates, i.e., if (1, 2) is such a genotype-pair, then so is (2, 1). This implies that *I*_2*a*_ = 0, as for every genotype pair (*a, b*) resulting in a non-zero 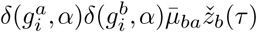, there always exist one other conjugate pair (*b, a*) which cancels this term through the 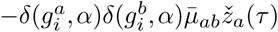 term.

Consider now *I*_2*b*_, and observe that the 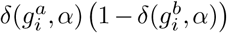 term is non-zero only when genotype *a*, but not genotype *b*, has allele *α* at locus *i*. Combining the above, we can thus write *I*_2_ as

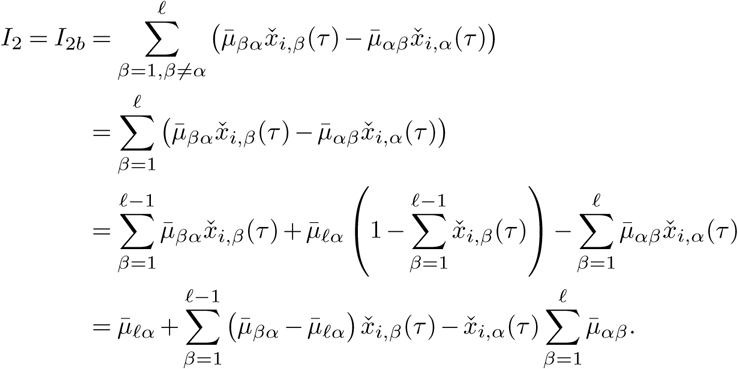

Note that in the third line above we have used the fact that the frequency of the reference allele is one minus the sum of the frequency of all other alleles.

Finally, we have

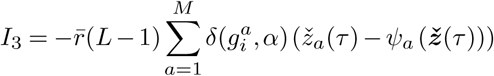

Similar to the biallelic case, we define

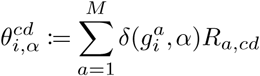

where 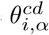 is the probability that genotypes *c* and *d* recombine to form a genotype which has an allele *α* at locus *i*. Next we split this summation into four summations, one for each of the four possible recombination scenarios. Namely, when both genotypes *c* and *d* have allele *α* at locus *i*, when both do not have allele *α* at locus *i* and when only of the genotypes has allele *α* at locus *i*. We thus have

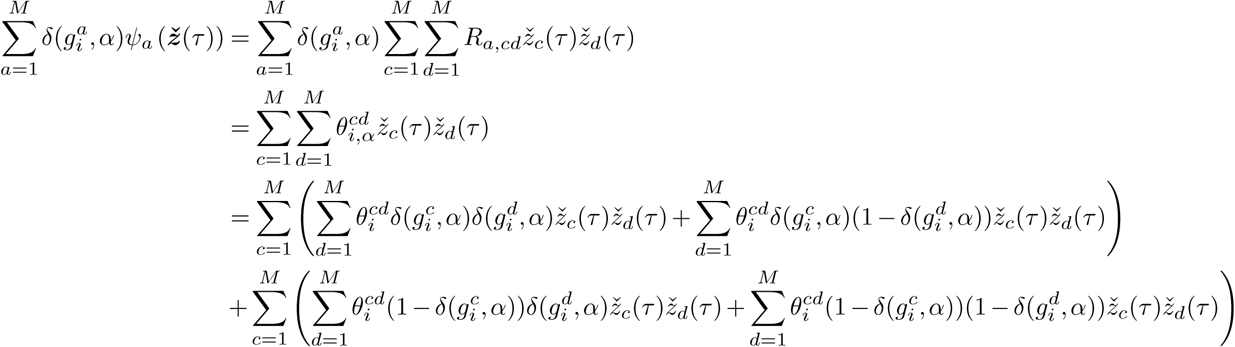

Moreover, noting that

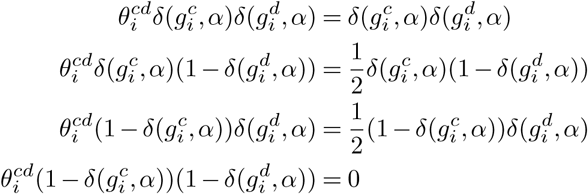

where the factor of 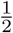 arises because there is a 50% chance that genotype *c* (*d*) with allele *v* at locus *i* and genotype *d* (*c*) which does not have allele *α* at locus *i* will recombine to a genotype with allele *α* at locus *i*, we have

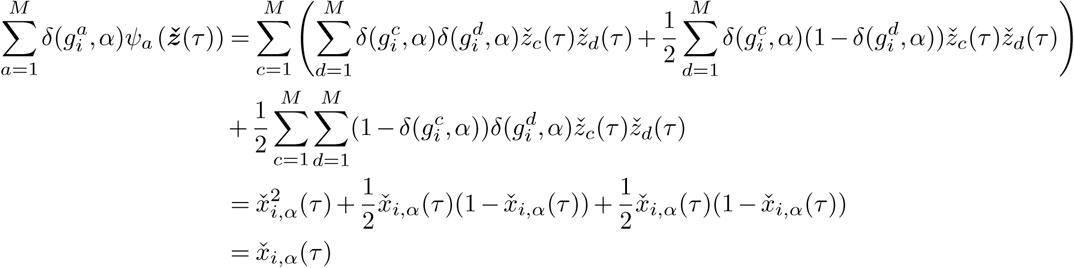

thus implying that 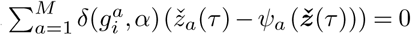 and hence *I*_3_ = 0.

The final expression for the drift vector is thus

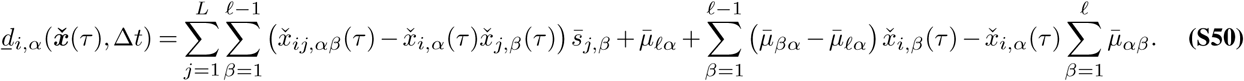

As expected, we observe that when there are two alleles, the mutation probability is symmetric and the reference allele is the WT, the expression in (S50) reduces to the expression in (S20).

We now move on to the diffusion matrix. This can be partitioned into *L*^2^ blocks each of size (ℓ− 1)× (ℓ − 1), where the (*i, j*)th block has (*α, β*)th entry

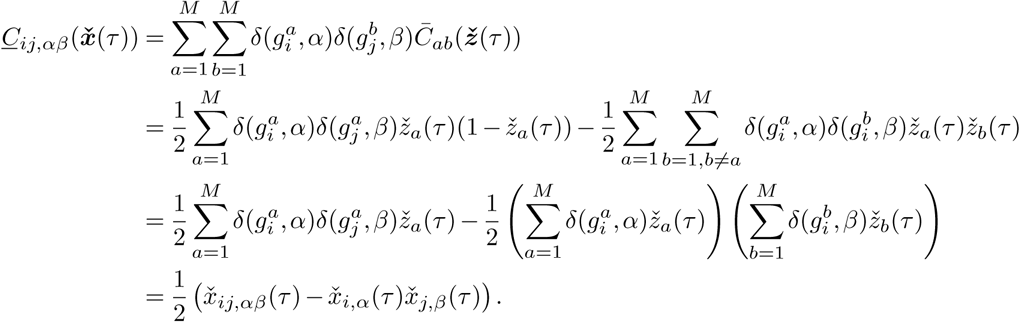

Following along the lines of the proof in Section 1.4, the probability of observing mutant allele frequencies (***x***(*t*_1_), ***x***(*t*_2_), *…*, ***x***(*t*_*K*_)), and hence the likelihood function (S42), is approximated by the path integral representation

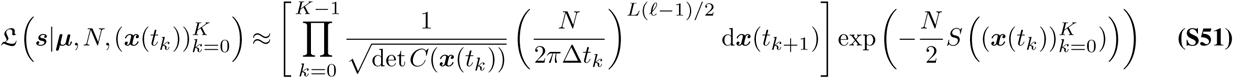

where 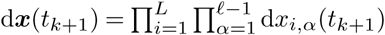 and

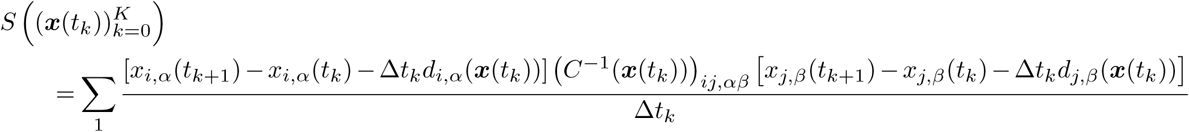

Here we have adopted the shorthand notation 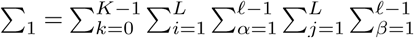, while (***X***)_*ij,αβ*_ indicates the (*α, β*)th entry of the (*i, j*)th block, with each block having dimension (ℓ − 1) × (ℓ − 1). Moreover, we have

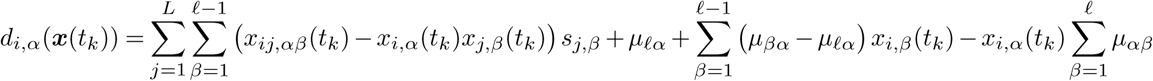

and

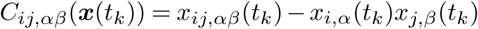

##### 2.5.3 The MPL estimator solution

Substituting the likelihood approximation (S51) and the prior

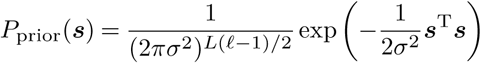

into the equivalent MAP problem

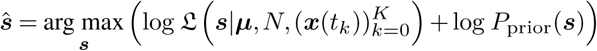

we take the vector derivative with respect to ***s*** and equate to zero. This yields the MPL estimate

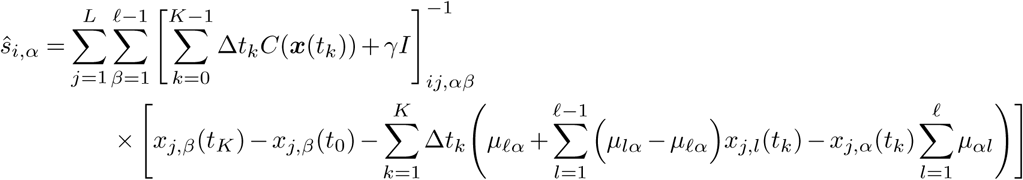

for *i* = 1, *…, L* and *α* = 1, *…, ℓ* − 1, which is the result quoted in (11) in Methods.

### 3. Data analysis

This section provides details about the synthetic data and its analysis used to obtain results presented in the paper.

#### 3.1. Simulated data generation and performance analysis

The results in Figure 1, Supplementary Figures 1 and 2 were obtained for a 50-locus biallelic system. We generated a population of *N* = 1000 sequences, each of length *L* = 50, and evolved the population according to the WF model with selection and mutation. We assumed loci to be biallelic with 0 representing the WT and 1 representing the mutant allele. Parameters for the simulations are included in the captions of Supplementary Figures 1 and 2. Scripts for generating and analyzing these data are located in the GitHub repository.

We compared MPL with seven methods from the literature: WFABC (3), FIT (4), ApproxWF (5), LLS (6), CLEAR (7), EandR (8), and IM (9). We compared the accuracy of these methods by measuring the AUROC for classification of beneficial and deleterious selection coefficients, as well as the normalized root-mean-squared error (NRMSE) of the inferred selection coefficients,

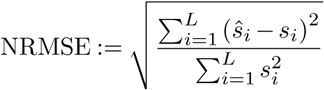

We also recorded the run time of all algorithms in the same computational environment. All values were averaged over samples from 100 WF simulations with the same underlying parameters.

#### 3.2. Implementation and data preprocessing specifications

We wrote custom codes for the FIT and IM methods, while for the remaining methods we used software implementations provided by the respective authors. Scripts used to generate the comparison results in the paper are available at the GitHub repository.

##### Preprocessing

Some methods required the input sequence or allele frequency data to be modified in order to produce reasonable results. This *pre-processing* was performed according to one or two rules, as indicated below. In describing these, we define a *trajectory* as a set of mutant allele frequencies at consecutively sampled time points. The start of a trajectory is marked by the first observation of a polymorphism, while the end is marked by either the fixation or loss of the mutant allele, or by the last sampled time point. Thus, over an entire observation period, it is possible to observe multiple “trajectories” at the same locus.

1. *Pre-filtering:* Trajectories were filtered such that (a) the minimum length of a trajectory was two time points and (b) the maximum (minimum) frequency was at least 0.05 (0.95) for a trajectory that started from a frequency of zero (one).
2. *Bit-flipping:* The definitions of WT and mutant were reversed at loci for which trajectories had initial frequency greater than 0.95.

###### MPL and SL

MPL and its single locus variant, the independent model described in the main text, were implemented in C++. No preprocessing of the sequence data was required for MPL and SL.

###### WFABC

Trajectories were converted to WFABC’s format using custom Matlab scripts and then analyzed using the code provided at http://jjensenlab.org/software (accessed on September 16, 2017). Initial tests revealed that we had to pre-process the raw sampled trajectories before applying the WFABC method in order to get meaningful results. All trajectories were pre-filtered except the ones that were rendered monomorphic due to pre-filtering. Bit-flipping was also applied. These processing steps removed spurious noisy blips from the trajectories and redefined WT/mutant, resulting in improved performance of the WFABC method.

###### ApproxWF

Trajectories were converted to an input format compatible with ApproxWF using custom Matlab scripts and then analyzed using the code provided at https://bitbucket.org/wegmannlab/approxwf/wiki/Home (commit fcc7964 dated: 2016-10-04 accessed on September 26, 2017). All trajectories were pre-filtered as outlined above, except the ones that were rendered monomorphic due to filtering. Bit-flipping was not used.

###### IM

We developed a custom MATLAB script to implement the IM method (9). Trajectories were pre-filtered, but not flipped. We used the filtering threshold of 0.1 (0.9) as specified by the authors of this method. The method was implemented according to the description given in ref. (9), which applies a simulated annealing algorithm to estimate the selection coefficients. We used the simulated annealing algorithm implementation provided in the Global Optimization Toolbox of MATLAB 2017a. We let the algorithm run for a maximum of 100, 000 iterations and stopped the algorithm when the average change in value of the objective function in the previous 10, 000 iterations was less than a tolerance value (default value of 10^−8^).

In all simulation runs, the optimization stopped in less than 100, 000 iterations, indicating that the algorithm had converged. To further validate our in-house implementation of IM, we applied it to data sets generated using the system parameters as in ref.(9) and under the same simulation setup; e.g., continuous long runs of several hundred thousand generations. The results were qualitatively similar to those reported in ref. (9). We found that IM’s procedure of calling trajectories (described in ref. (9)), was sensitive to the evolutionary parameters used, e.g., population size, mutation probability (*N* = 10, 000 and *µ* = 5 × 10^−7^ were used in ref. (9)), as well as to the simulation condition of not allowing a locus to mutate after it has reached fixation (in the simulation setup of IM, a locus that reaches fixation remains at fixation for the next 3200 generations). For the simulation results presented in this paper, where *N* = 1000, *µ* = 10^−4^ and there are no restrictions on mutations at a locus reaching fixation, we had to change the parameters of the procedure for calling trajectories so that the algorithm returned meaningful results. Using the default parameters of ref. (9) for calling trajectories resulted in the IM method missing the start/end of trajectories. The modified parameters are specified in Matlab scripts in the GitHub repository.

###### LLS

Trajectories were fed directly into the LLS code provided by the authors at https://github.com/ThomasTaus/poolSeq. A small number of loci resulted in a “N/A” selection coefficient estimate, which occurred when the loci had a frequency of zero or one for the vast majority of time points, and having a frequency close to zero or one for the other (small number of) time points. These loci were excluded from the analysis.

###### FIT

No additional preprocessing was required. The trajectories were analyzed using custom Matlab scripts.

###### CLEAR

Simulation data was converted into a format readable by CLEAR using custom Python scripts. No preprocessing of the data was necessary.

###### EandR

Simulation data was converted into a format readable by EandR using custom Python scripts. No preprocessing of the data was required.

